# Thrombospondin-1 Promotes Circuit-Specific Synapse Formation via β1-Integrin

**DOI:** 10.1101/866590

**Authors:** Sehwon Koh, Suva Roy, Oznur Eroglu, Samuel Strader, William J. Chen, Jeremy N. Kay, Greg D. Field, Cagla Eroglu

## Abstract

Glial cells regulate synaptic connectivity during development, but whether they selectively instruct the formation of specific synaptic circuits is not known. Here we show that the major perisynaptic glia of the retina, the Muller glia (MG), control the proper establishment of the direction-selective (DS) circuit by a synaptogenic protein, Thrombospondin 1 (TSP1). We found that TSP1 promotes excitatory synapse formation specifically in <u>o</u>n-<u>o</u>ff <u>D</u>irection-<u>S</u>elective retinal <u>G</u>anglion <u>C</u>ells (ooDSGCs). Lack of TSP1 leads to reduced synapse formation within the inner plexiform sublayers containing DS-circuit, resulting in deficits of ooDSGC function. Even though pan-TSP receptor, α2δ-1, interaction is required for TSP1-induced synapse formation, the ooDSGC-subtype specificity of TSP1 is conferred by a second neuronal TSP1 receptor, β1-Integrin. Furthermore, conditional deletion of β1-Integrin in ooDSGCs results in diminished excitatory synapse formation without disturbing laminar organization showing that MG-secreted TSP1 controls circuit-specific synapse formation via β1-Integrin.

## INTRODUCTION

Neural circuit function is dependent on the precision and specificity of synaptic connectivity. Correspondingly, aberrant formation or loss of synapses results in circuit dysfunction which is a hallmark of many neurological disorders. Therefore, elucidating the molecular and cellular underpinnings of how specific synaptic connections are established and maintained is of fundamental importance for understanding central nervous system (CNS) function in health and disease.

Most studies of selective synapse formation have focused on cell-cell recognition between neurons. Several families of neuronal cell adhesion molecules and secreted proteins have been shown to play important roles in synapse specificity (Berns et al., 2018; Chih et al., 2006; Duan et al., 2014; Krishnaswamy et al., 2015; Tran et al., 2009; Zipursky and Sanes, 2010). However, it remains unclear whether the specificity of synapse formation can be entirely explained by an adhesion code among circuit partners, or if other cellular and molecular mechanisms also play an instructive role.

Glial cells have been shown to be strong regulators of synapse formation, maturation and plasticity (Allen and Eroglu, 2017; Araque et al., 2014; Khakh and Sofroniew, 2015; Ma et al., 2016; Ullian et al., 2004; Wu et al., 2015). In the brain, astrocytes are the principal perisynaptic glial cells and they strongly stimulate excitatory synapse formation through secretion of the synaptogenic molecules such as Thrombospondin family proteins (TSPs) (Christopherson et al., 2005; Risher and Eroglu, 2012). All five members of the TSP family proteins (TSP1-5) induce an increase in the number of synapses made between neurons in glia-free cultures by interacting with their common neuronal receptor, the calcium channel subunit α2δ-1 (Christopherson et al., 2005; Eroglu et al., 2009; Risher et al., 2018).

The ability of astrocytes to promote synapse formation, via the use of a variety of synaptogenic factors like TSPs, raises the possibility that astrocytes induce selective synapse formation between particular neuronal partners. If this were the case, astrocytes would need to release these synaptogenic cues at locations where those synapses should form, and/or neurons belonging to specific circuits should specifically respond to distinct astrocytic factors. Whether such selectivity in synaptogenic astrocyte-neuron signaling occurs is unclear. This lack of clarity is primarily due to the extremely complicated architecture of astrocyte-neuron interactions and the immense heterogeneity of neurons and synapses in the brain.

To investigate the potential roles of astroglia in formation of specific circuits, we utilized the retina, in which the laminar architecture of the synaptic layers facilitates identification of synapses belonging to particular circuits. The inner plexiform layer (IPL) neuropil contains at least five sublayers, each hosting synapses that process distinct aspects of the visual scene (Lefebvre et al., 2015; Sanes and Yamagata, 2009). Developing retinal neurons express a variety of cell-cell recognition molecules that guide their projections to particular IPL sublayers, where they form synapses with their circuit partners (Duan et al., 2018; Kay et al., 2012; Matsuoka et al., 2011; Peng et al., 2017; Sun et al., 2013; Yamagata and Sanes, 2008). However, how glia-neuron interactions contribute to the establishment and function of the specific retinal circuitry is largely unexplored.

The retina contains a specialized perisynaptic astroglial cell type, Muller glia (MG), which produce TSP1 and TSP2 during postnatal development and well into adulthood (Koh et al., 2018). TSP2 is enriched in the outer plexiform layer (OPL) containing photoreceptor synapses, whereas TSP1 is enriched in two distinct sublayers of the inner plexiform layer (IPL) (Koh et al., 2018) that host synapses between starburst amacrine cells (SACs), bipolar cells and on-off direction-selective retinal ganglion cells (ooDSGCs): the core elements of the direction selective (DS) circuit (Kay et al., 2011; Mauss et al., 2017; Wei and Feller, 2011). The shared neuronal receptor for TSPs, α2δ-1, is highly enriched in both synaptic layers of the retina (Koh et al., 2018). Thus, the retina provides a powerful model to study how glia-neuron interactions, mediated by TSPs, shape the formation and maturation of distinct neuronal circuits.

In this study, we identified a molecular mechanism through which TSP1 selectively controls DS circuit assembly and function in the retina. Using a combination of purified rat retinal neuron cultures and mouse genetics, we found that TSP1 promotes excitatory synapse formation specifically onto ooDSGCs; whereas its close homolog, TSP2 does not exhibit this specificity. We also found that lack of TSP1 leads to impaired synapse development specifically in DS circuit IPL layers, causing deficits in direction tuning among ooDSGCs. Furthermore, even though both TSP1 and TSP2 require their common receptor, α2δ-1, for their synaptogenic activity, another neuronal TSP1 receptor, β1-Integrin, confers the cell-type specific synaptogenic effect of TSP1. Accordingly, conditional ablation of β1-Integrin in ooDSGCs sharply reduces the number of excitatory synapses made onto their dendrites. Taken together, these results provide novel molecular insights into how MG shape DS circuit development and how TSP1/β1-Integrin-signaling directs synapse formation in a cell-type specific manner.

## RESULTS

### Thrombospondin-1 promotes synapse formation onto a specific subpopulation of retinal ganglion cells

To mechanistically examine the role of TSPs in retinal synapse formation, we used purified rat retinal ganglion cells (RGCs) in culture, which have been extensively utilized to discover the molecular underpinnings of astrocyte-induced synapse formation (Barres et al., 1988; Christopherson et al., 2005; Ullian et al., 2001). RGCs, which are purified from P7 rat retinas, form very few synapses when cultured alone in a serum-free growth media of known composition, but addition of astrocyte or retinal glial feeder layers or astroglia-conditioned media robustly increases the number of excitatory synapses (Christopherson et al., 2005; Koh et al., 2018). The synapses formed on the RGCs are identified by immunohistochemical methods by quantifying the colocalization of pre- (Bassoon, green) and post- (Homer, red) synaptic markers. The pre- and post-synaptic proteins are in distinct neuronal compartments (axons and dendrites, respectively) and would only appear to co-localize at synapses, wherein they are extremely close contact (Ippolito and Eroglu, 2010).

To test whether TSP1 and TSP2 differentially affect formation of synapses between RGCs, we treated them with recombinant human TSP1 or TSP2 (150 ng/mL) for 6 days (DIV4 to DIV10), and quantified the synapses formed onto RGCs (Figure 1A). As shown before (Christopherson et al., 2005; Eroglu et al., 2009; Koh et al., 2015), TSP1 and TSP2 both robustly increased synapse numbers in RGCs compared to cells treated with culture growth media only (GM Only-Control) (Figures 1B and 1C, TSP1, 2.35±0.284-fold, TSP2, 3.05±0.42-fold, One-way ANOVA, p<0.0001). The synaptogenic effects of TSP1 and TSP2 were not statistically different (Figure 1C). To determine if there were differences in the response of individual RGCs to TSP1 versus TSP2 treatments, we plotted the fold increase in synapse number per cell (population density plot of relative fold increase normalized to the mean of GM Only-Control) in all three conditions and found that only a subgroup of RGCs responded to TSP1 treatment strongly, while others did not (Figure 1D, orange, arrow versus arrowhead). On the other hand, TSP2 uniformly increased the synapse numbers in all RGCs (Figure 1D, blue, i.e. right shifted bell curve). These observations indicate that TSP2 induces synapse formation in most of the RGCs, whereas TSP1 exerts strong synaptogenic effects only onto a subset of RGCs. These in vitro results, together with our previous finding that TSP1 is enriched in two sublayers of the IPL containing the DS-circuit specific synapses (Koh et al., 2018), suggested that TSP1 may control formation of synapses specifically onto the ooDSGCs.

**Figure 1.**
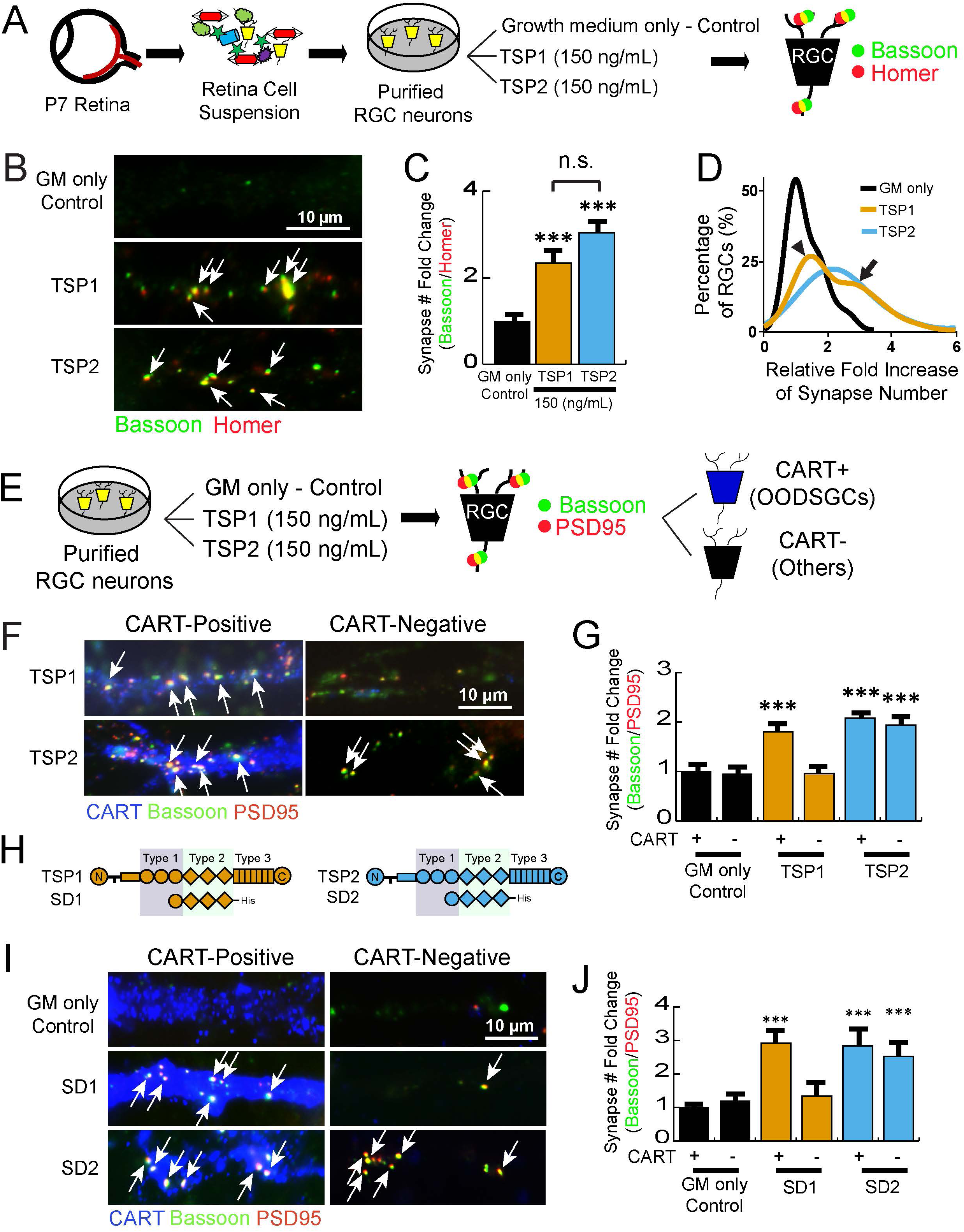
TSP1 promotes synapse formation in CART-positive subpopulation of RGCs. **(A)** Schematic representation of the experimental design. RGCs are isolated from P7 rat retinas through a series of immunopanning. The RGCs are treated with growth medium only (<u>G</u>rowth <u>M</u>edia Only-Control) or recombinant human TSP1 or TSP2 (150 ng/mL) for 6 days. Synapses are identified by the co-localization of pre-(Bassoon, green) and post- (Homer, red) synaptic markers. **(B)** Representative images of the synapses formed on the RGCs treated with either TSP1 or TSP2. Synapses, co-localized synaptic puncta (yellow), are marked with white arrows. Full images of the RGCs are available in the supplement. **(C)** Quantification of fold increase in the number of co-localized synaptic puncta demonstrates both TSP1 and TSP2 induces synapse formation in RGCs (n=30 cells/condition, One-way ANOVA, *** p<0.0001). Fold increase is calculated by normalizing the number of synapses per cell with the number of synapses per cell in GM Only-Control. **(D)** Density plot showing the distribution of RGCs by the fold increase of synapse number. TSP1 (orange), but not TSP2 (blue), induces synapse formation in a subpopulation of RGCs (marked by arrow compare to arrowhead). **(E)** Schematic representation of the experimental design. After 6-days treatment of TSP1 or TSP2, On-Off direction selective RGCs (OODSGCs) are labeled by a marker, CART (blue). Synapses are visualized by the co-localization of synaptic markers, Bassoon (pre-, green) and PSD95 (post-, red). **(F)** Representative images of the synapses formed on the CART-positive or -negative RGCs treated with either TSP1 or TSP2. Full images of the RGCs are available in the supplement. **(G)** TSP1 induces synapse formation in the CART-positive RGCs while TSP2 promotes synapse formation in both CART-positive and – negative RGCs (n=30 cells/condition, One-way ANOVA, *** p<0.0001). **(H)** The domain structures of full-length TSP1, TSP2 and synaptogenic domain fragments of TSP1 (SD1) or TSP2 (SD2). **(I)** Representative images of the synapses formed on the CART-positive or -negative RGCs treated with either SD1 (150 ng/mL) or SD2 (150 ng/mL). Full images of the RGCs are available in the supplement. **(J)** Treatment of the synaptogenic domain fragments of TSP1 (SD1) or TSP2 (SD2) demonstrated identical synaptogenic function to the full-length TSPs (n=30 cells/condition, One-way ANOVA, p<0.0001). All data were expressed as mean±SEM.

To test this possibility, we repeated the *in vitro* RGC neuron culture assay shown in Figure 1A, but this time we included an ooDSGCs marker, <u>C</u>ocaine and <u>A</u>mphetamine <u>R</u>egulated <u>T</u>ranscript (CART) (Kay et al., 2011). After 6 days of TSP1 or TSP2 treatment, ooDSGCs were labeled by anti-CART, along with the synaptic markers (Bassoon (pre) and PSD95 (post)) (Figure 1E). Quantification of the synapse number in CART-positive versus negative RGC populations showed that TSP1 specifically induces excitatory synapse formation onto the CART-positive, but not CART-negative RGCs (Figure 1F-G, Figure S1A-B). On the contrary, TSP2 promoted synapse formation onto both CART-positive and CART-negative RGCs (Figure 1F-G, Figure S1A-B). The gabapentin receptor (also known as calcium channel subunit) α2δ-1 is a common receptor for TSPs that is required for the synaptogenic functions of all TSPs, including TSP1 and TSP2 (Eroglu et al., 2009). In agreement with this, gabapentin (32 µM), the anti-epileptic anti-analgesic drug that inhibits the interaction between TSPs and α2δ-1 (Eroglu et al., 2009), blocked both TSP1 and TSP2-induced synapse formation in RGC cultures, regardless of the RGC subtype (Figure S1C-D). This result shows that α2δ-1 interaction is required for both TSP1 and TSP2-mediated synaptogenesis.

TSP1 and TSP2 are large trimeric multi-domain proteins with high homology. The domains of TSP1 and TSP2 that are crucial for their synaptogenic functions were previously identified (Eroglu et al., 2009). The fragment of TSP1 or TSP2 composed of one Type I (properdin-like) repeat and three of the Type II (EGF-like) repeats (Figure 1H) have previously been shown to be as synaptogenic as the full-length TSPs (Eroglu et al., 2009). To determine if the synaptogenic domain of TSP1 (SD1) also induces RGC subtype-specific synapse formation, SD1 and the corresponding synaptogenic domain of TSP2 (SD2) were expressed in human embryonic kidney (HEK) cells and purified from the conditioned medium (Figure S1C, see methods for details). RGC cultures were treated with purified SD1 and SD2 (150 ng/mL) for 6 days (Figure S1D). Subsequent synapse quantification revealed that, similar to their corresponding full-length TSPs, SD1 only promoted synapse formation onto CART-positive RGCs, whereas SD2 induced synapse formation onto both CART-positive and CART-negative RGCs (Figure 1I-J). Taken together, these results show that TSP1 promotes synapse formation specifically in CART-positive ooDSGCs, and the synaptogenic fragment of TSP1 is sufficient for its cell type-selective synaptogenic activity.

### Lack of TSP1 decreases the number of a specific subset of IPL synapses in vivo

The IPL of the retina is grossly divided into five synaptic sublayers (S1-S5, Figure 2A). TSP1 was previously found to be enriched in two of these sublayers, S2 and S4 in P30 retinas (Koh et al 2018). These synaptic sublayers harbor the DS circuit connections made between cholinergic starburst amacrine cells (SACs), ooDSGCs, and subtypes of bipolar neurons that arborize to S2 (Off), S4 (On), or both layers (Figure 2A). Previous studies have shown that the dendrites of SACs and ooDSGCs are closely intertwined (Wei and Feller, 2011). Thus, staining with the SAC specific markers, such as <u>CH</u>oline <u>A</u>cetyl<u>T</u>ransferase (CHAT) marks the S2 and S4 sublayers and the excitatory synapses within these synaptic layers can be identified as the colocalization of VGluT1 (pre-synaptic, green) and PSD95 (post-synaptic, red) (Figure 2B).

**Figure 2.**
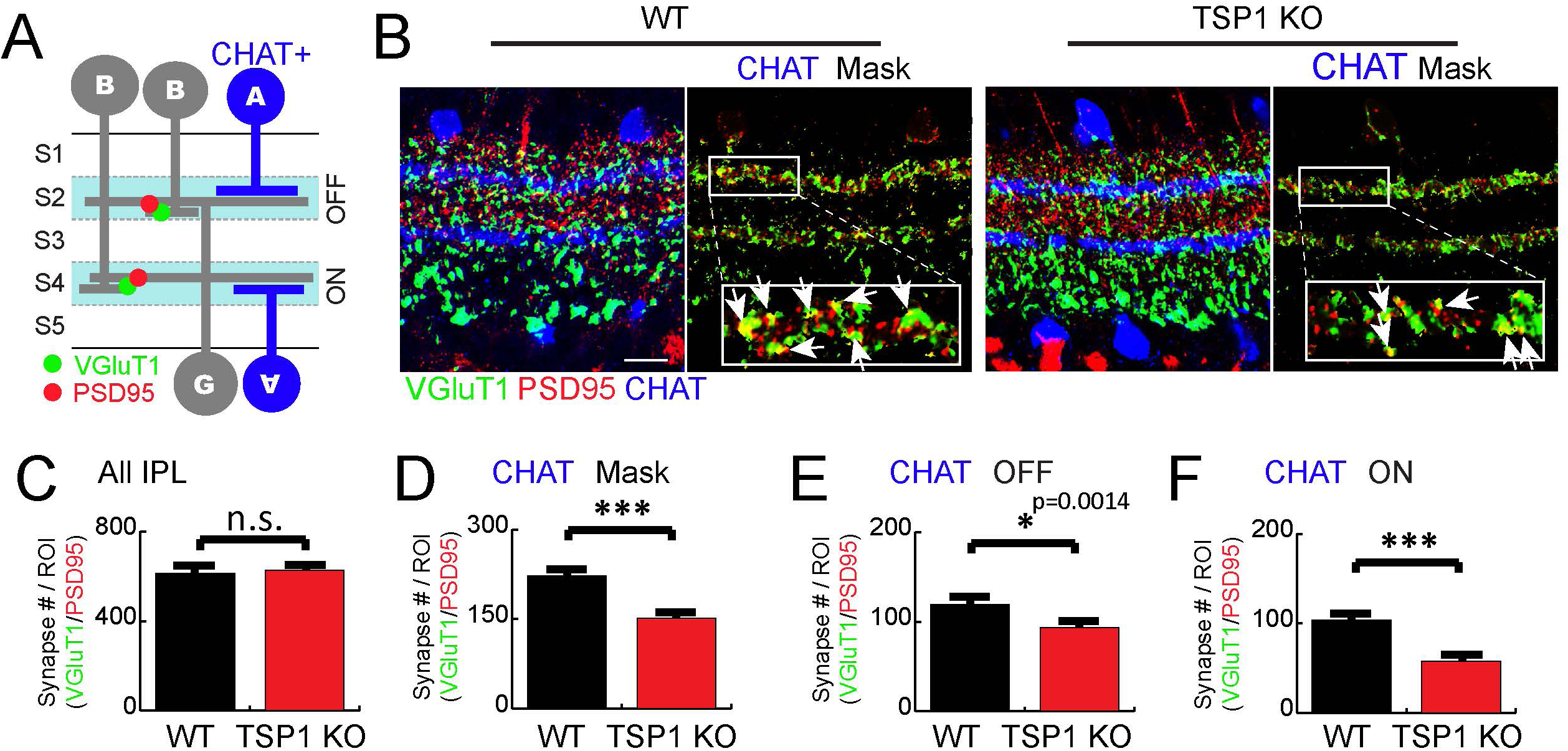
TSP1 is required for the excitatory synapse development of the DS circuit IPL sublayers. **(A)** Schematic of IPL and cell types composing the DS circuit IPL sublayers. CHAT-positive starburst amacrine cells (blue) project to two IPL sublayers (off and on). The excitatory synapses formed between bipolar cells and ganglion cells are identified by the co-localization of synaptic markers, VGluT1 (pre-, green) and PSD95 (post-, red). **(B)** Representative images of the excitatory synapses in WT and TSP1 KO retinas at P30. DS circuit IPL sublayers are visualized by a starburst amacrine cells specific antibody, CHAT (blue) and excitatory synapses are labeled by VGluT1 (pre-, green) and PSD95 (post-, red). Only the synapses within two sublayers (S2 and S4) where ooDSGCs project are shown by masking with CHAT staining (right panels). The inlets (white boxes) are shown in higher magnification and the co-localized synaptic puncta (merge, yellow) are marked with white arrows. **(C)** There was no significant change in the number of excitatory synapses formed when all IPL sublayers are examined (n=3 mice per genotype, t-test, n.s. non-significant). **(D)** Quantification of excitatory synapses specifically formed in CHAT-layers demonstrates significant reduction in synapse number in TSP1 KO retinas (n=3 mice per genotype, t-test, *** p<0.0001). Both **(E)** Off- and **(F)** On-layers showed significantly reduced synapse number in TSP1 KO (n=3 mice per genotype, t-test, *** p<0.0001). (n=3 mice per genotype, t-test, *** p<0.0001).

To test if TSP1 is required for excitatory synapse development in the IPL, we quantitatively analyzed the number of VGluT1/PSD95 positive synapses using established methods (see materials and methods) in TSP1 knock-out (TSP1 KO) animals at P30, an age during which TSP1 protein is abundant in the IPL (Koh et al., 2018). The retinas from littermate wild-type (WT) animals were used as controls. When we quantified the number of VGluT1/PSD95 positive excitatory synapse densities within the entire IPL, we found no significant differences between WT and TSP1 KOs (Figure 2B-C).

To determine if the lack of TSP1 specifically affected the number of excitatory synapses formed in the S2 and S4 sublayers, these sublayers were marked by co-immunostaining with CHAT (Figure 2B). To quantify synapses located exclusively within the DS sublayers, CHAT signal was used to generate a mask and synapses outside the CHAT mask were excluded from the quantification (Figure 2B, right panels). Quantification of excitatory synapses within these CHAT-positive layers showed a significant reduction in the number of VGluT1/PDSD95 positive synapses in TSP1 KOs compared to littermate WT controls (Figure 2D). The synapse numbers were reduced in both S2 (off) and S4 (on) layers of the TSP1 KOs compared to WTs (Figure 2E-F).

Because the inhibitory synapses formed between SACs and ooDSGCs are crucial components that govern direction selectivity of the DS-circuit (Poleg-Polsky et al., 2018), we next investigated whether the lack of TSP1 affected inhibitory synapse numbers in the retina. The inhibitory synapses were identified as the colocalization of Bassoon (pre-synaptic, green) and Gephyrin (post-synaptic, red), and another SACs marker, <u>V</u>esicular <u>A</u>cetyl<u>Ch</u>oline <u>T</u>ransporter (VAChT), was utilized to distinguish DS sublayers (Figures 3A-B). Interestingly, we found a significant reduction in inhibitory synapse numbers throughout the IPL (Figure 3C) and within the DS circuit layers (Figure 3D). Both S2 (off) and S4 (on) sublayers were affected in TSP1 KOs (Figures 3E-F).

**Figure 3.**
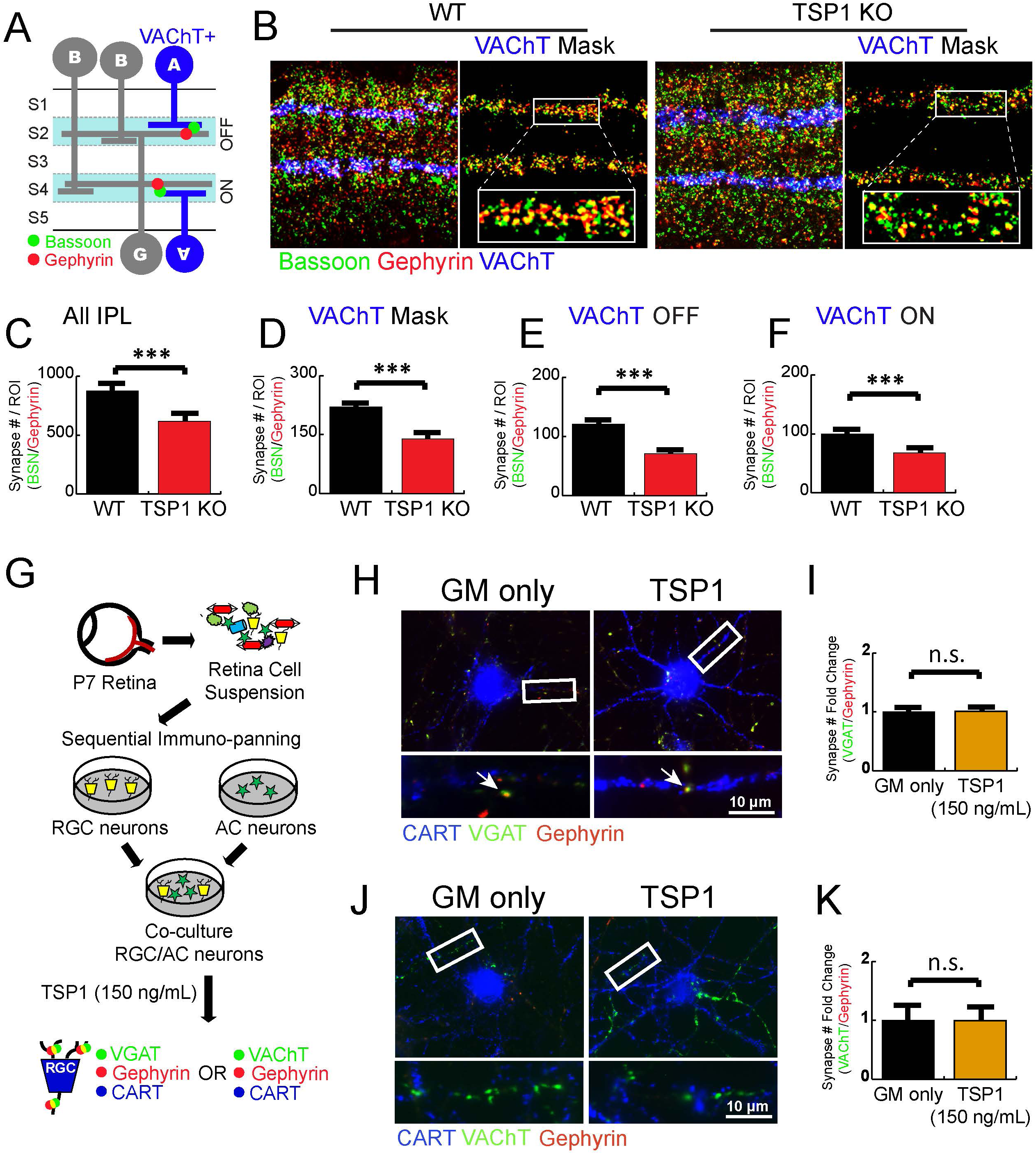
Lack of TSP1 leads to reduced inhibitory synapse number in the IPL. **(A)** Schematic of IPL and cell types composing the DS circuit IPL sublayers. CHAT-positive starburst amacrine cells (blue) project to two IPL sublayers (off and on). The inhibitory synapses formed between starburst amacrine cells and retinal ganglion cells are labeled by synaptic markers, Bassoon (pre-, green) and Gephyrin (post-, red). **(B)** Representative images of the inhibitory synapses and CHAT-positive dendrites in WT and TSP1 KO retinas at P30 (left panels). The synapses within two sublayers (S2 and S4) where ooDSGCs project are shown by masking with CHAT (right panels). The inlets (white boxes) are shown in higher magnification. **(C)** Number of inhibitory synapse within the IPL is significantly reduced in TSP1 KO compared to WT (n=3 mice per genotype, t-test, *** p<0.0001). **(D)** Quantification of inhibitory synapses formed in the two sublayers demonstrates significant reduction in TSP1 KO retinas (n=3 mice per genotype, t-test, *** p<0.0001). Both **(E)** Off- and **(F)** On-layers showed significantly reduced inhibitory synapse number in TSP1 KO (n=3 mice per genotype, t-test, *** p<0.0001). **(G)** Schematic representation of the experimental design. RGCs and ACs are isolated from P7 rat retinas through a series of immunopanning, then co-cultured. The co-cultured RGCs/ACs are treated with <u>G</u>rowth <u>M</u>edia Only-Control or recombinant human TSP1 (150 ng/mL) for 6 days. GABAergic or Cholinergic synapses formed on ooDSGCs (CART-positive, blue) are identified by the co-localization of pre- (GABAergic, VGAT, green) and post- (Gephyrin, red) or pre- (Cholinergic, VAChT, green) and post- (Gephyrin, red) synaptic markers. Representative images of the **(H)** GABAergic and **(J)** Cholingergic synapses formed on the RGCs treated with either growth medium (GM Only-Control) or TSP1. Quantification of fold increase in the number of co-localized synaptic puncta demonstrates that TSP1 do not increase either **(I)** GABAergic or **(K)** Cholinergic synapse numbers. (n=30 cells/condition, Student’s t-test, n.s., not significant). Fold increase is calculated by normalizing the number of synapses per cell with the number of synapses per cell in GM Only-Control.

These results suggested two possibilities: 1) TSP1 also controls the formation of inhibitory synapses in the retina, or 2) the number of inhibitory synapses are reduced in TSP1 KO retinas as a compensatory mechanism due to the reduction in the excitatory synapses. To test the first possibility, we tested whether TSP1 is sufficient to induce inhibitory synapse formation between retinal neurons. To do so, we utilized an *in vitro* co-culture of amacrine cells (AC) with RGCs. The ACs and RGC neurons were isolated by a series of immuno-panning steps as previously described (Barres et al., 1988; Goldberg et al., 2002). The RGCs and ACs were then cultured together at 1:4 ratio (15K RGCs and 60K ACs respectively), mimicking *in vivo* cell proportions, and supplemented with full-length TSP1 or cultured in growth media alone (GM Only-Control) for 6 days (Figure 3G). The numbers of GABAergic (<u>V</u>esicular <u>GA</u>BA <u>T</u>ransporter, VGAT/Gephyrin) and Cholinergic (VAChT/Gephyrin) synapses formed onto CART-positive RGCs were assessed (Figure 3G). The results showed that TSP1 is not sufficient to induce either GABAergic (Figures 3H-I) or cholinergic (Figure 3J-K) synapse formation. Collectively, our *in vitro* and *in vivo* results show that TSP1 is required for the proper formation of excitatory synapses onto the ooDSGCs. Our results also indicate that proper TSP1 signaling within the retina, possibly due to its role in wiring excitatory synapses, is also critical for inhibitory synapse development in the IPL.

### TSP1 overexpression promotes synapse formation in DS circuit IPL sublayers

Thbs1, mRNA that transcribes TSP1 (Figure S2A) is highly enriched in Glutamine Synthetase positive (GS, green, Figure S2A) MG cell bodies (Figure S2A-B, also see (Koh et al., 2018)), suggesting that MG are the primary source of TSP1 in the IPL. Thus, we next investigated whether the re-expression of TSP1 in the MG is sufficient to stimulate excitatory synapse formation within the DS circuit. To reinstate TSP1-signaling in MG, we overexpressed (OE) the synaptogenic domain of TSP1 (SD1) using MG-specific AAVs (Figure 4A) and drove its expression under the control of gfaABC1D promoter (Lee et al., 2008). To identify the cells that express SD1, a fluorescence reporter gene, mCherry was also expressed under the same promoter through an internal ribosome entry site (IRES). A viral construct with the same backbone that only expresses mCherry was used as the control virus (Figure 4B) and each mouse were injected in left eye with SD1-expressing virus and the contralateral eyeball of the same animal was used for control AAV (Figure 4A-B).

**Figure 4.**
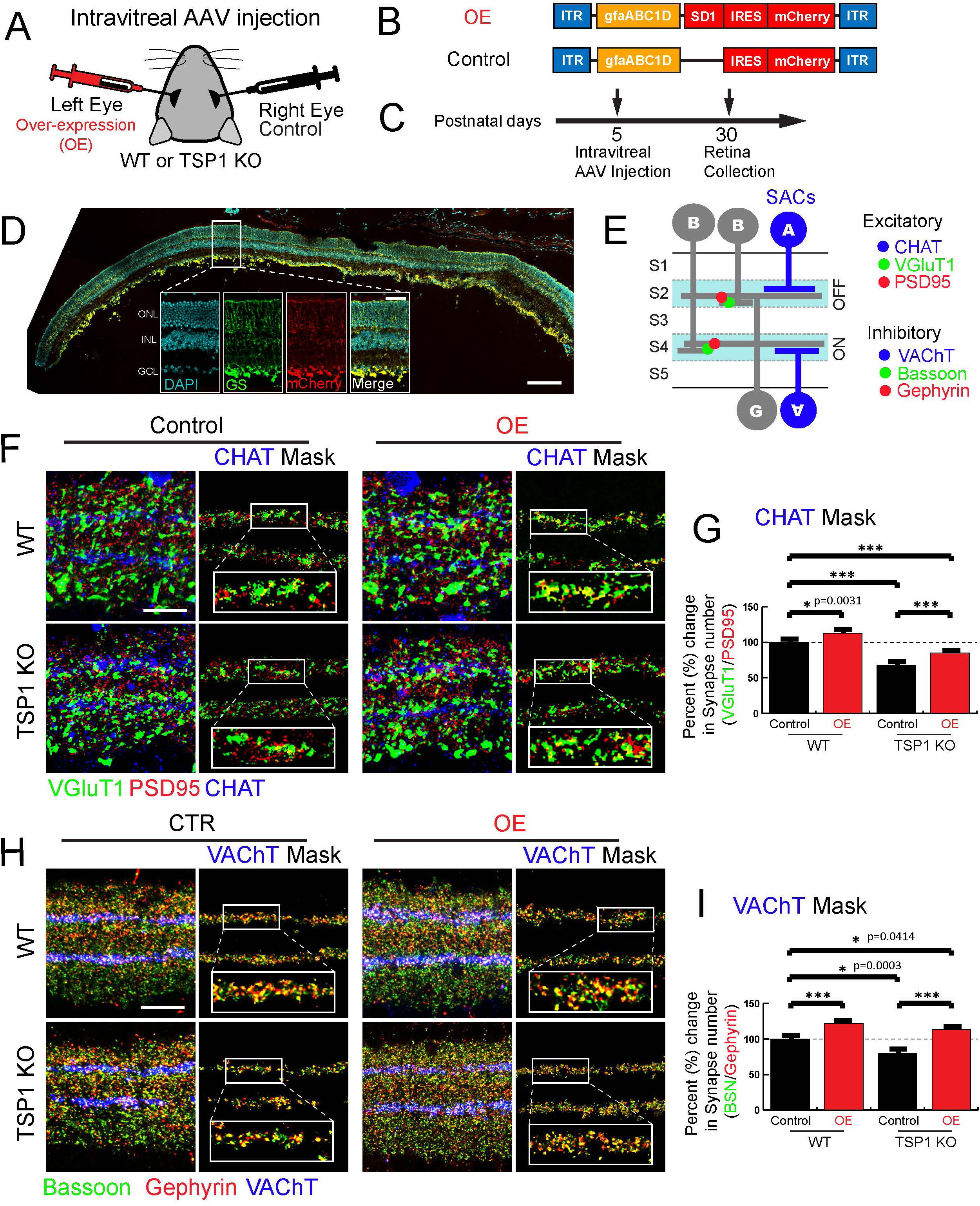
MG-produced TSP1 is sufficient to induce synapse formation in DS circuit IPL sublayers. **(A)** AAV particles that express SD1-IRES-mCherry (over-expression, left eye) or mCherry only (control, right eye) are intravitreally injected at P5. **(B)** Schematics of DNA constructs used for AAV production. Expression of SD1-IRES-mCherry (for over-expression) and mCherry alone (control) is driven by glia-specific gfaABC1D promoter. **(C)** Intravitreal injection of AAV particles is performed at P5, then retinas are collected at P30. **(D)** Co-localization of Muller-glia specific marker, GS (green), and mCherry (red) demonstrates Muller-glia specific expression of the transgene. **(E)** Schematic of IPL and cell types composing the DS circuit. CHAT (or VAChT)-positive SACs (blue) project to two IPL sublayers (S2 and S4). The excitatory synapses formed between bipolar cells and ganglion cells are labeled by synaptic markers, VGluT1 (pre-, green) and PSD95 (post-, red). The inhibitory synapses are labeled by Bassoon (pre-, green) and Gephyrin (post-, red). **(F)** Representative images of the excitatory synapses in WT and TSP1 KO retinas with over-expression of SD1 at P30. The inlets (white boxes) are shown in higher magnification and the co-localized synaptic puncta (merge, yellow) are marked with white arrows. Over-expression (OE) of SD1 induced excitatory synapse formation within the DS circuit sublayers in both wild-type and TSP1 KO **(F and G)** (n=3-4 mice per genotype per treatment, One-way ANOVA, *** p<0.0001). Relative percent change is calculated by normalizing the number of synapses to littermate WT retinas. **(H and I)** Representative images of the inhibitory synapses in WT and TSP1 KO retinas with over-expression of SD1 at P30. The inlets (white boxes) are shown in higher magnification and the co-localized synaptic puncta (merge, yellow) are marked with white arrows. Over-expression (OE) of SD1 induced inhibitory synapse formation within the DS circuit sublayers in both wild-type and TSP1 KO **(H and I)** (n=3-4 mice per genotype per treatment, One-way ANOVA, *** p<0.0001). Relative percent change is calculated by normalizing the number of synapses to WT.

The AAV particles were intravitreally delivered at P5, and then the retinas were collected at P30 (Figure 4C). Successful transduction of the AAV in the MG was confirmed by the co-localization of the MG marker (GS, green) and mCherry (red) (Figure 4D). Excitatory synapses (VGluT1/PSD95) and inhibitory synapses (Bassoon/Gephyrin) located within the DS circuit sublayers were identified as described above using the SAC marker, CHAT or VAChT (Figure 4E). Synapse quantification revealed that overexpression of SD1 in MG is sufficient to promote excitatory synapse formation in both WT and TSP1 KO retinas (Figure 4F-G). Overexpression of SD1 in WT retinas led to a significant increase in excitatory synapse numbers in the DS circuit sublayers compared to control virus-injected retinas (Figure 4G). Importantly, re-expression of SD1 in TSP1 KO, significantly recovered the synapses formed in the DS circuit sublayers. Likewise, number of inhibitory synapses formed within the DS circuit sublayers were significantly increased in both WT and TSP1 KO (Figure 4H-I). Overexpression of SD1 in MG also increased excitatory and inhibitory synapse numbers in non-DS IPL regions (Figure S2C-D), suggesting that localized and controlled secretion of TSP1 at S2 and S4 may be necessary for spatially restricting the synaptogenic effects of TSP1. In summary, these results show that MG-mediated TSP1-signaling is necessary and sufficient for controlling excitatory and inhibitory synaptic development in the IPL.

### Lack of TSP1 impairs fine direction tuning function of ooDSGCs

Next, to examine how lack of TSP1 impacts DS circuit function, we measured the direction selective responses of ooDSGCs in TSP1 KO retinas. Using a large-scale multielectrode array (MEA), we recorded spikes from hundreds of RGCs simultaneously from ex-vivo retinas (Litke, 2004; Yao et al., 2018). DSGCs were identified and distinguished from other RGCs and further sub-classified into ON and ON-OFF subtypes based on their responses to bars moving in twelve different directions at two different speeds (see Methods, Figure 5A-B) (Vaney et al., 2012; Yao et al., 2018). The fraction of DSGCs to the total number of RGCs recorded was similar between WT (∼8%) and TSP1 KO (∼7%) retinas (Figure 5A), indicating the overall number of direction selective RGCs did not differ significantly between the genotypes. The ooDSGCs in both WT and TSP1 KO retinas exhibited a cruciform organization, with each cell exhibiting a preferred direction aligned to one of four cardinal directions (Figure 5D). However, the direction tuning curves of ooDSGCs were significantly wider in the TSP1 KOs compared to WT retinas (Figure 5B-C), indicating that direction tuning of ooDSGCs is altered in TSP1 KOs.

**Figure 5.**
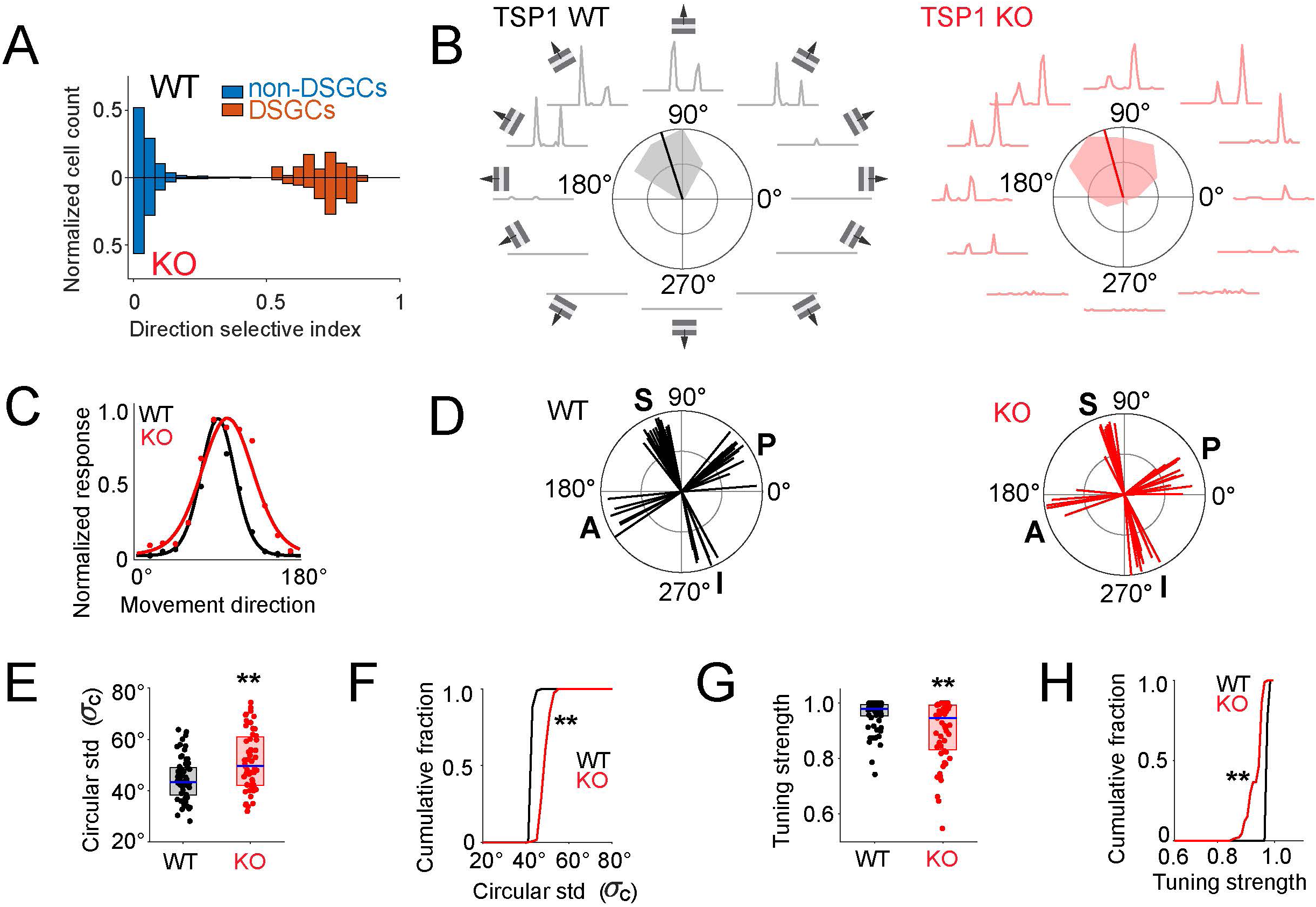
Lack of TSP1 leads to deficits in DS tuning of ooDSGCs. **(A)** Distribution of direction selective index across all RGCs recorded in the multielectrode array. The RGCs with DSI above 0.5 are classified as DSGCs in WT (top, n=70/754) and in TSP1 KO (bottom, n=64/777). The scale on Y axis to the right (orange) is magnified to highlight the distribution of DSGCs. **(B)** Peri-stimulus time histograms (PSTHs) of representative ooDSGCs from WT (black) and TSP1 KO (red) retinas. PSTHs represent response averaged over 8 repeats of a bar moving in 12 different directions. Normalized response for each direction is shown by the contour and the preferred direction is shown as a line from the center, in the polar plots. **(C)** Representative direction tuning curves from WT (black) and KO (red) retinas. **(D)** Preferred directions of each ooDSGC recorded from retinas of a representative pair of WT and TSP1 KO animals. The direction and length of each line represent the preferred direction and strength of population normalized response respectively. The four ooDSGC subtypes are annotated as S: Superior, I: Inferior, A: Anterior, P: Posterior. **(E)** Circular standard deviation (σc) of individual ooDSGCs is increased in TSP1 KO (red) compared to the WT (black) (Paired Kolmogorov-Smirnov test, ** asymptotic p<0.01, WT: n=65, KO: n=55). The population median, 0.25 quartile and 0.75 quartile of σc are shown by blue line, lower and upper limits of the shaded box respectively. **(F)** Cumulative distribution of circular standard deviation demonstrates higher population mean circular standard deviation of ooDSGCs in TSP1 KO (red) compared to the WT (black). **(G)** Tuning strength (normalized differential response to preferred direction (PD) versus null direction (ND)) of individual ooDSGCs is decreased in TSP1 KO (red) compared to the WT (black). The population median, 0.25 quartile and 0.75 quartile of tuning strength are shown by blue line, lower and upper limits of the shaded box respectively. **(H)** Cumulative distribution of tuning strength shows lower population mean tuning strength of ooDSGCs in TSP1 KO (red) compared to the WT (black). (Population size of ooDSGCs in KO: n=55 and WT: n=65, from 2 mice of each genotype. Paired Kolmogorov-Smirnov test, ** asymptotic p<0.01)

To assess the change in direction tuning of ooDSGCs in TSP1 KO compared to WT retinas, we quantified two aspects of direction tuning: (1) the tuning width, which was quantified by the circular standard deviation (σ*_c_*), and (2) the tuning strength, which was quantified by the difference between the preferred and null direction responses normalized by their sum (see Methods). The tuning of ooDSGCs was significantly broader in TSP1 KO retinas than WT (Figures 5E-F; median and median absolute deviation of σ*_c_* = 52°±11° for TSP1 KO and 43°±8° for WT, ** p<0.01). This difference in tuning width emerged clearly in the cumulative distribution plot of the circular standard (Figure 5F, ** p<0.01). Furthermore, the tuning strength of ooDSGCs in TSP1 KO retinas was reduced compared to WT retinas (0.90±0.09 for TSP1 KO and 0.97±0.04 for WT) (Figures 5G-H, ** p<0.01). Collectively, these results show that lack of TSP1 results in diminished direction selectivity across the population of ooDSGCs. Thus, TSP1-mediated synapse formation in the direction-selective circuit is required for establishing precise DS-tuning among ooDSGCs.

### Third type I repeat of TSP1 is required and sufficient for its cell-type specific synaptogenic activity

How does TSP1 induce synapse formation in an ooDSGC-specific manner? To answer this question, we first set to identify the region within the synaptogenic fragment of TSP1 (SD1) that is responsible for conferring cell-type specificity. To do so, we took advantage of the highly-conserved domain structure and organization of TSP1 and TSP2 and generated SD1 and SD2 chimeras (Figure 6A). First, we took the SD2 fragment of TSP2, which induces excitatory synapse formation onto all RGCs (Figure 1I-J), and swapped in either the third type 1 (properdin-like) repeat and/or the third type 2 (EGF-like) repeat from TSP1 (Figures 6B-C). We chose to test the involvement of the third type 1 or type 2 domains of TSP1 in cell-type specificity, because these regions of TSP1 are important domains for receptor interactions (Calzada et al., 2004; Eroglu et al., 2009; Resovi et al., 2014). The chimeric fragments were expressed in HEK cells and were purified from culture media and analyzed with SDS-PAGE to confirm their purity (Figure 6C). Of note, even though the SD1 and SD2 are composed of the same number of amino acids, the molecular weight of SD2 is slightly higher than SD1 due to different glycosylation status (Hoffmann et al., 2012). Swapping either the third type I or type II repeat from TSP1 onto SD2 had no effect on molecular weight, indicating that the glycosylation sites that are specific to SD2 are located within the first and/or second type II repeats (Figure 6C).

**Figure 6.**
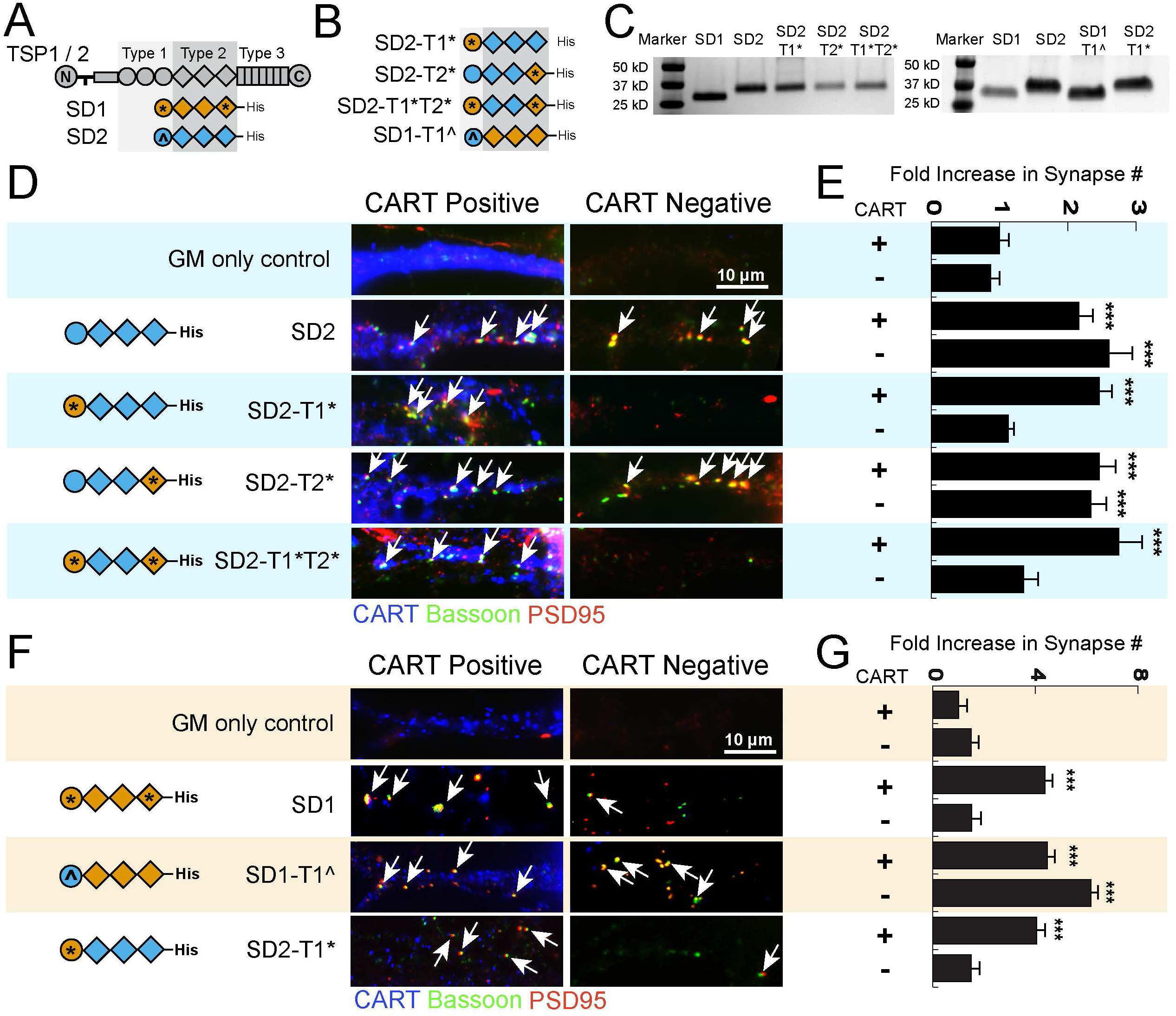
The third Type 1 repeat of TSP1 confers RGC subtype specificity. **(A)** Schematic representation of synaptogenic domain (SD) structures of the TSP1 and TSP2. **(B)** Schematic representation of the purified chimeric synaptogenic domains. Synaptogenic fragment of TSP1 (SD1), TSP2 (SD2) or the chimeric SD1 and SD2 fragments were treated to the RGCs isolated from P7 rat retinas at 150 ng/mL concentration. **(C)** SDS-PAGE analysis of purified fragments. **(D)** Representative images of the synapses formed on the RGCs treated with various synaptogenic fragments. Synapses (white arrows) are identified as co-localized synaptic puncta (yellow) of pre- (Bassoon, green) and post-synaptic (PSD95, red) markers. ooDSGCs are marked by CART (blue). Full images of RGCs are available in the supplementary figure 3. **(E)** Quantification of fold increase in the number of co-localized synaptic puncta (n=30 cells/condition) demonstrates that replacement of type 1 repeat (SD2-T1*), but not type 2 repeat (SD2-T2*), convert synaptogenic function of SD2. SD2-T1* induce synapse formation specifically in CART-positive RGCs. Fold increase is calculated by normalizing the number of synapses per cell with the number of synapses per cell in CART+ RGCs alone (n=30 cells/condition, One-way ANOVA, *** p<0.0001). **(F)** Representative images of the synapses formed on the RGCs treated with different fragments. RGCs isolated from P7 rat retinas are treated with purified synaptogenic fragment of TSP1 (SD1) or the chimeric SD1 fragments that contains type1 repeat of TSP2 (SD1-T1^) or chimeric SD2 fragments containing type1 repeat of TSP1 (SD2-T1*) at 150 ng/mL concentration. Synapses, co-localized synaptic puncta (yellow) of pre- (Bassoon, green) and post- synaptic (PSD95, red) markers, are marked with white arrows. CART (blue) visualizes ooDSGCs. Full images of RGCs are available in the supplementary figure 3. **(G)** Quantification of fold increase in the number of co-localized synaptic puncta demonstrates that chimeric SD1 containing Type 1 repeat of TSP2 (SD1-T1^) induces synapse formation in both CART-positive and –negative RGCs mimicking synaptogenic function of TSP2. Fold increase is calculated by normalizing the number of synapses per cell with the number of synapses per cell in CART+ RGCs alone (n=30 cells/condition, One-way ANOVA, *** p<0.0001).

Next, we tested the synaptogenic activity of these chimeras *in vitro* using purified RGCs (as described above, Figure 1). We found that the chimera (SD2-T1*) generated by switching only the third type 1 repeat of TSP1 onto SD2 induces synapse formation specifically in CART-positive RGCs. Whereas SD2-T2* induced synapse formation onto both CART-positive and –negative RGCs similar to SD2 itself (Figures 6D-E and S3A). The chimeric SD2 fragment that contains both type I and type II repeats of TSP1 (SD2-T1*T2*) also displayed cell-type specificity in synaptogenic activity by only inducing synaptogenesis onto CART-positive cells. Taken together these results show that the third type 1 repeat of TSP1 is sufficient to confer cell type-specific synaptogenic activity to SD2 (Figures 6D-E and S3A).

To test the necessity of the third type I repeat of TSP1 to confer cell type-specificity, we produced another chimera by replacing the third type I repeat in SD1 with the same repeat from TSP2 (SD1-T1^) (Figure 6B). The molecular weights were similar between the native SD1 and the chimera SD1-T1^ (Figure 6C). The resulting chimera SD1-T1^ was highly synaptogenic and its synaptogenic effect was not limited to CART-positive cells like SD1, but rather affected both CART-positive and CART-negative RGCs like SD2 (Figures 6F-G and S3B). Taken together, these results show that the third type I repeat of TSP1 is both necessary and sufficient to confer the CART-positive RGC-type specificity for its synaptogenic function. These findings also suggest that the cell-type specific effects of TSP1 in synaptogenesis is mediated through an interaction between the third type1 repeat of TSP1 and a neuronal TSP1 receptor that confers this specificity.

### β1-Integrin is the neuronal TSP1 receptor that confers cell-type specificity for synaptogenic activity

A considerable number of cell surface receptors interact with TSPs in general and TSP1 in particular (Resovi et al., 2014). The shared synaptogenic receptor for TSPs, α2δ-1, is highly enriched all IPL sublayers of the retina (Koh et al., 2018) and Cacna2d1 mRNA transcript encoding for this protein is expressed in similar amounts across all RGCs (Figure S4A). β1-Integrin is another known TSP1 receptor which interacts with the third type I repeat of TSP1 (Calzada et al., 2004) suggesting that TSP1/β1-Integrin interaction may play a role in conferring RGC-subtype specificity. To determine if the β1-Integrin is expressed in the RGCs in general and DSGCs in particular, we checked mRNA expression of Itgb1, that transcribes β1-Integrin, in previously reported gene expression profiles composed of 13 retinal neuron subtypes (Kay et al., 2012). Interestingly, Itgb1 expression was highest in RGCs (Thy1^+^), and enriched specifically in Hb9^+^, CART^+^ and Drd4^+^ RGC subgroups that include ooDSGCs (Figure S4B). To confirm these gene profiling results, we utilized quantitative RNA imaging analysis (i.e. RNAscope) based on fluorescence in situ hybridization (FISH), and we determined that in the ganglion cell layer (GCL) of the retina, Itgb1 is almost exclusively expressed by RGCs compared to the other cell types (Figure S4C-D). Importantly, Itgb1 expression is significantly enriched in CART-positive ooDSGCs compared to CART-negative RGCs (Figure S4E). These observations suggested β1-Integrin as the TSP1 receptor that is required for the specificity of TSP1 to induce synapse formation only in CART-positive RGCs.

To test this possibility, purified rat RGCs were treated with full-length TSP1 or TSP2 for 6 days in the presence of either a function-blocking antibody against β1-Integrin or a serotype-matched control IgG (Figure 7A). The control IgG had no effect on the synaptogenic activity of either TSP (Figures 7B-C and S5A-B); however, in agreement with a role for β1-Integrin interactions on TSP1-induced synapse formation, the function blocking β1-Integrin antibody abolished TSP1-mediated synapse formation onto CART-positive cells (Figure 7B-C). TSP2 still significantly enhanced synapse formation onto both CART-positive and CART-negative RGCs even in the presence of the function-blocking antibody against β1-Integrin (Figure. 7B-C, Figure S5A-B). Taken together these results show that β1-Integrin function is required for TSP1-mediated synapse formation onto CART-positive RGCs.

**Figure 7.**
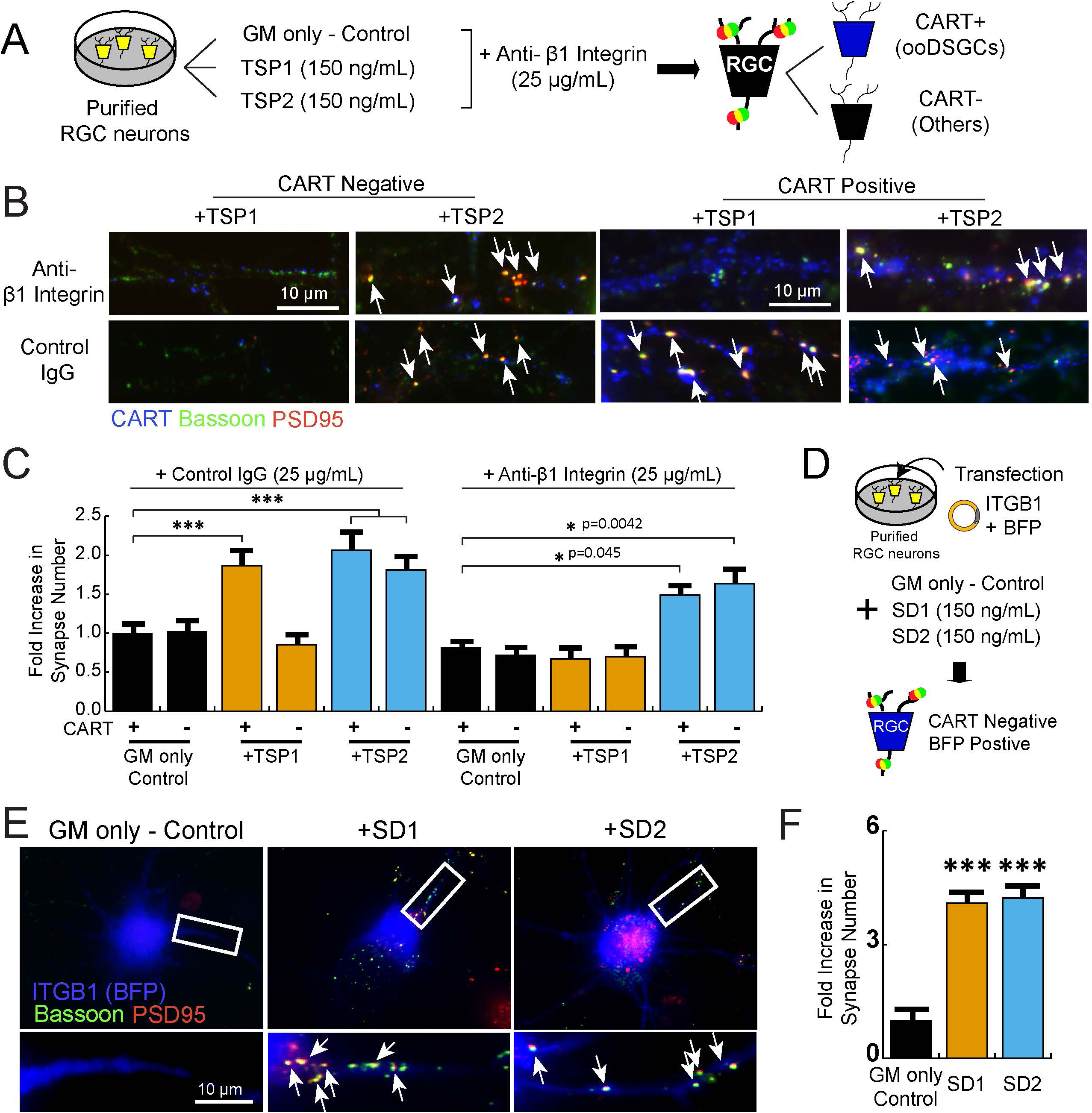
ooDSGC-specific synaptogenic activity of TSP1 is mediated via β1-Integrin. **(A)** Schematic representation of the experimental design. Purified RGCs isolated from P7 rat retinas are treated with recombinant human TSP1 or TSP2 (150 ng/mL) in the presence of the function blocking antibody against Integrin β1 or control IgG. RGCs were labeled with a ooDSGC marker, CART (blue), and synaptic markers, Bassoon (pre-, green) and PSD95 (post-, red). **(B)** Representative images of the synapses formed on the RGCs treated with either full length TSP1 or TSP2 in the presence of integrin β1 function blocking antibody or control IgG. Synapses, co-localized synaptic puncta (yellow) of pre- (Bassoon, green) and post-synaptic (PSD95, red) markers, are marked with white arrows. CART (blue) visualizes ooDSGCs. Full images of RGCs are available in the supplementary figure. Quantification of fold increase in the number of co-localized synaptic puncta from the RGCs treated with **(C)** function blocking antibody or control IgG demonstrates blocking Integrin β1 specifically inhibits TSP1-mediated synaptogenic activity but not TSP2. Fold increase is calculated by normalizing the number of synapses per cell with the number of synapses per cell in CART+ RGCs in GM Only-Control condition (n=30 cells/condition, One-way ANOVA, *** p<0.0001). **(D)** Schematic representation of the experimental design. Purified RGCs isolated from P7 rat retinas are transfected with the DNA plasmids transcribing ITGB1 and BFP, then treated with synaptogenic fragments, SD1 or SD2 (150 ng/mL). Then the synapses formed onto the RGCs expressing BFP but negative for CART were imaged for synapse quantification. **(E)** Representative images of the synapses formed on the RGCs expressing ITGB1 and BFP. The RGCs are treated with either SD1 or SD2, then synapses (white arrows) are labeled as co-localized puncta (yellow) of pre- (Bassoon, green) and post-synaptic (PSD95, red) markers. **(F)** Quantification of fold increase in the number of co-localized synaptic puncta demonstrates that expression of ITGB1 sufficient for TSP1-mediated synapse formation in CART-negative RGCs. (n=20-30 cells/condition, One-way ANOVA, *** p<0.0001).

Next we tested whether increasing the expression of β1-Integrin is sufficient to enable TSP1 to induce synapse formation onto CART-negative RGCs. DIV 5 RGCs were transfected with a plasmid containing ITGB1 cDNA, to express human β1-Integrin, concurrently with blue fluorescence protein (BFP) reporter. Two days post transfection the RGCs were supplemented with either SD1 or SD2 and cultured for 6 additional days (Figure 7D) and synapses were immunolabeled and quantified as described before. We found that the overexpression of ITGB1 is sufficient for SD1 to induce synapse formation in CART-negative RGCs (Figure 7E-F). This result shows that increasing β1-Integrin abundance in CART-negative RGCs is sufficient to prime these neurons to respond to TSP1. Taken together, these results strongly indicate that β1-Integrin expression and function is sufficient and required to confer the RGC subtype-specific synaptogenic activity of TSP1.

### β1-Integrin is required in ooDSGCs for circuit-specific excitatory synapse development in vivo

To determine if β1-Integrin is required for synapse formation in the DS circuit *in vivo*, we conditionally knocked-out (cKO) the Itgb1 gene encoding for β1-Integrin from ooDSGCs in the developing mouse retina. To do so, the transgenic mice with a floxed allele of Itgb1 (Itgb1 flox, (Raghavan et al., 2000)) were crossed with mice expressing CreER under the Cadherin 6 promoter (Cdh6-CreER) that was previously demonstrated to drive Cre expression in the ooDSGCs upon Tamoxifen injection (Kay et al., 2011). Because β1-Integrin plays important roles in cell migration and neurite outgrowth and sublayer-specific projection and elaboration (Neugebauer and Reichardt, 1991; Pasterkamp et al., 2003; Santos et al., 2012), we chose to knockout β1-Integrin after P9-P10 to allow ooDSGCs undergo normal early neuronal development and maturation, but ablate β1-Integrin right before the period of active synapse formation in the IPL (∼P14) (Fisher, 1979). The Cre-mediated recombination event was monitored by a Cre reporter allele, Ai14, (Rosa-CAG-loxP-STOP-loxP-tdTomato) (Figure 8A). All the transgenic mice we used for the quantification analyses carried one allele of tdTomato (Tg/0) and one allele of Cre (Tg/0). The Cre-expression was induced by tamoxifen (20 mg/mL, subcutaneous) injections at P9 and P10. Retinas from β1-Integrin cKO mice (Itgb1 f/f, Cdh6-CreER (Tg/0), tdTomato (Tg/0)) and their littermate controls (Itgb1 +/+; Cdh6-CreER (Tg/0); tdTomato (Tg/0)) were collected at P30 (Figure 8B).

**Figure 8.**
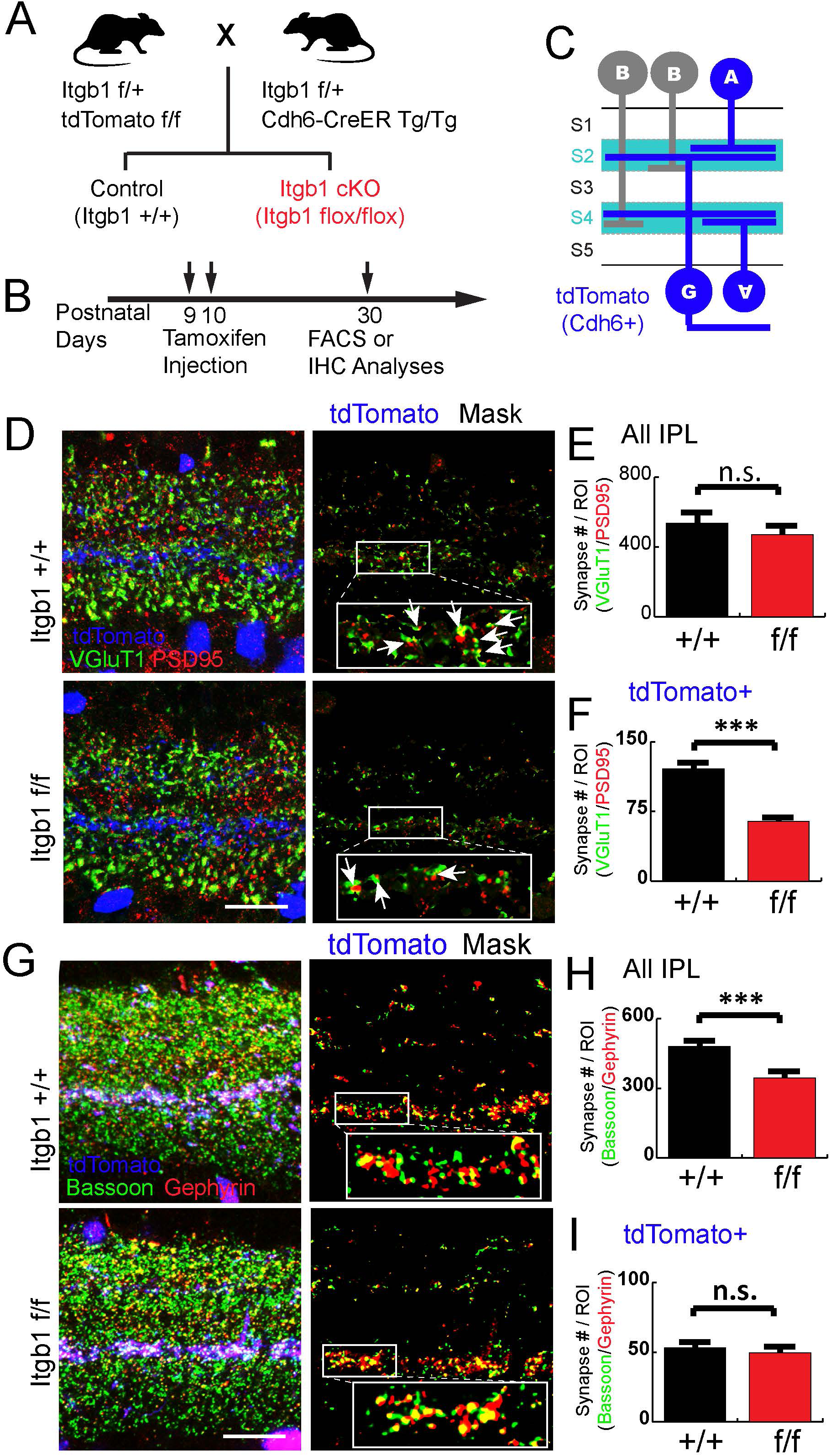
β1-Integrin is required for the DS circuit synapse development. **(A)** Schematic representation of the transgenic mice breeding strategy. Retinas from the littermate wild-type (WT, Itgb1 +/+) and Integrin β1 cKO (Itgb1 f/f) were collected at P30. **(B)** Tamoxifen (20 mg/mL) was subcutaneously delivered twice at P9 and P10 (0.6 mg each day) then retinas were collected at P30. **(C)** Schematic of IPL and cell types that express Cadherin 6 (blue). **(D)** Representative images of the excitatory synapses in WT and Itgb1 cKO retinas at P30. Cadherin6-positive DS circuit IPL sublayers are visualized by tdTomato (blue, left panels) and excitatory synapses are labeled by VGluT1 (pre-, green) and PSD95 (post-, red). Only the excitatory synapses within tdTomato positive dendrites are shown by masking with tdTomato staining (right panels). The inlets (white boxes) are shown in higher magnification and the co-localized synaptic puncta (merge, yellow) are marked with white arrows. **(E)** There is no significant change in the number of excitatory synapses within the IPL between control (+/+) and cKO (f/f). **(F)** Quantification of excitatory synapses within tdTomato-positive dendrites demonstrates significant reduction in synapse number in Itgb1 cKO retinas (n=3 animals per genotype, t-test, *** p<0.0001). **(G)** Representative images of the inhibitory synapses in WT and Itgb1 cKO retinas at P30. The inhibitory synapses are labeled by Bassoon (pre-, green) and Gephyrin (post-, red)). Only the synapses within tdTomato positive dendrites are shown by masking with tdTomato staining. The inlets (white boxes) are shown in higher magnification and the co-localized synaptic puncta (merge, yellow) are marked with white arrows. **(H)** There is a significant reduction in the number of inhibitory synapses within the IPL between control (+/+) and cKO (f/f) (n=3 animals per genotype, t-test, *** p<0.0001). **(I)** Quantification of inhibitory synapses within tdTomato-positive dendrites demonstrates that there is no change in inhibitory synapse number between Itgb1 WT and cKO retinas (n=3 animals per genotype, t-test, n.s., not significant)

To confirm the loss of Itgb1 transcript in Cre expressing retinal cells in cKOs, P30 retinas were collected and enzymatically dissociated into a single cell suspension. The tdTomato-positive vs –negative cells were isolated by fluorescence-activated cell sorting (FACS) from both control and cKO retinas. Total mRNA was isolated from FACS-sorted cells followed by cDNA synthesis (Figure S6A). A total of 0.4% FACS-sorted cells were tdTomato-positive in both control and cKO suggesting that the ablation of Itgb1 after P9-P10 period did not alter total number of Cdh6-CreER expressing cells in the retina (Figure S6B). Expression level of Itgb1 mRNA was measured by quantitative real-time polymerase chain reaction (qRT-PCR) using the cDNA isolated from the tdTomato-positive or –negative cells. We found about a 70% reduction in Itgb1 mRNA expression in cKO cells compared to controls; whereas, there was no change in Itgb1 expression in tdTomato-negative cells from both genotypes (Figure S6C).

Previous studies have shown that Cdh6 is specifically enriched in both ooDSGCs and SACs (Kay et al., 2011), suggesting that Cre-expression under the control of Cdh6 promoter would target these DS-circuit cells. In agreement with this, the Cre-expressing tdTomato-positive neurons were located in the INL (SACs) and GCL (both SACs and RGCs) (Figures 8C and S6D). As expected, the processes of the Cdh6-CreER-positive cells project to the CHAT-positive S2 and S4 sublayers of the IPL, indicating that Cre is expressed specifically in SACs and ooDSGCs (Figures 8D and S6D). Interestingly, a higher number of Cre-positive dendrites were observed on the S4 (ON) layer compared to S2 (OFF) (Figure S7D) in both control and cKO. Importantly, elimination of β1-Integrin in ooDSGCs after P9-P10 did not alter neurite outgrowth or dendritic projections as shown by comparable distribution of Cre-positive neurites along the IPL layers between controls and cKOs (Figure S6D-E). Similarly, cKO of β1-Integrin in SACs had no effect in dendritic arborization of the CHAT-positive processes to S2 and S4 (Figure S6D-E). Taken together, these results show that Cdh6-CreER after P9-P10 specifically targets SACs and ooDSGCs in the retina and elimination of β1-Integrin by this manipulation does not alter neurite outgrowth or layer distribution. Because β1-Integrin is not highly expressed in non-RGCs that include SACs in the GCL (Figure S4C-F), we concluded that Cdh6-CreER line is suitable for DS circuit-specific deletion of β1-Integrin to interrogate its function in synaptogenesis onto ooDSGCs in vivo.

Next, we tested whether conditional ablation of β1-Integrin in ooDSGCs affects excitatory synapse development. To do so, we stained retinal sections from cKO and control mice with VGluT1/PSD95 to mark synapses between BCs and RGCs (Figure 8D). We found no significant changes in the number of excitatory synapses when all IPL sublayers were analyzed (Figure 8E). However, when we analyzed only the synapses that were associated with Cre-positive dendrites by using a mask with tdTomato signal that was overlaid on top of VGluT1/PSD staining (Figure 7D, right panels), we found that excitatory synapses were greatly reduced in β1-Integrin cKO dendrites compared to littermate controls (Figure 8F).

In parallel, we also quantified the number of inhibitory (Bassoon/Gephyrin) synapses that were made onto the tdTomato-positive dendrites in β1-Integrin cKO and control retinas (Figure 8G-I). There were no significant differences in the number of Bassoon/Gephyrin synapses between β1-Integrin cKO dendrites compared to controls (Figure 8H). However, when we quantified the total number of inhibitory synapses within all IPL sublayers, we found a significant reduction in the number of inhibitory synapses in β1-Integrin cKOs compared to controls (Figure 8H). This result shows that ablation of β1-Integrin in SACs and ooDSGCs does not directly impact the number of inhibitory synapses onto their dendrites in IPL. However, these results suggest that the deletion of β1-Integrin in SACs and ooDSGCs reduction is sufficient to cause a reduction in inhibitory synaptic connectivity throughout the IPL.

Taken together our results show that β1-Integrin is required for excitatory synapse development in the DS circuit. The expression of β1-Integrin is enriched in ooDSGCs, and cKO of β1-Integrin in ooDSGCs results in a significant reduction in the number of excitatory synapses in the retinal sublayers that host the DS-circuit.

## DISCUSSION

In this study we took advantage of the highly organized structure and well characterized function of retinal inner plexiform synaptic layers and determined that glia-secreted TSP1 specifically promotes the formation of excitatory synapses within the retinal circuit that mediates the direction selectivity of visual responses. Our findings showed that synaptogenic activity of TSP1 is RGC subtype specific and we found that MG-secreted TSP1 is necessary and sufficient to promote excitatory synaptogenesis onto ooDSGCs. Consequently, in the absence of TSP1, the DS-circuit development is hampered and the direction-tuning functions of ooDSGCs are impaired.

Importantly, we also identified the molecular mechanism that confers the cell-type specificity of TSP1’s synaptogenic activity. This is achieved through the third Type I (properdin-like) repeat of TSP1 which interacts with β1-Integrin (Calzada et al., 2004). β1-Integrin is enriched in ooDSGCs. Blocking β1-Integrin function inhibits TSP1-induced synapse formation onto the ooDSGCs in vitro. Furthermore, overexpressing β1-Integrin in RGCs other than ooDSGCs elicits a synaptogenic response in these cells upon TSP1 treatment. Additionally, a conditional knock-out of β1-Integrin in ooDSGCs *in vivo* reduces excitatory synapse formation onto their dendrites. Thus, TSP1 instructs excitatory synapse development in a DS circuit-specific manner via β1-Integrin. More generally, these findings also demonstrate that different TSP isoforms play non-redundant roles in controlling synaptic circuit assembly in the CNS and provides novel insights into how glial cells shape synaptogenesis in specific neural circuits via secretion of soluble factors.

### TSP1 induces circuit-specific excitatory synaptogenesis through a crosstalk between β1-Integrin and α2δ-1-signaling

TSPs are oligomeric extracellular matrix glycoproteins that interact with multiple cell surface molecules and secreted ligands to control diverse and important roles in cell adhesion, tissue remodeling and morphogenesis (Bornstein and Sage, 2002; Lawler, 2002; Resovi et al., 2014). TSP1-4 are found in the brain and are produced primarily by astrocytes (Cahoy et al., 2008). All five TSPs are synaptogenic and their synaptogenic function is processed through their common receptor, the calcium channel subunit α2δ-1, that signals through the actin cytoskeleton modulator, Rac1 (Eroglu et al., 2009; Risher et al., 2018).

Here, we found that even though α2δ-1, is required for both TSP1 and TSP2-mediated synapse formation, it is not sufficient for TSP1-induced synaptogenesis in all RGCs. TSP1 can only effectively induce synapse formation onto ooDSGCs and its synaptogenic function is dependent on both α2δ-1 and β1-Integrin function. Previously, TSP1 or TSP2 fragments that contain the Type II EGF-like repeats were shown to be synaptogenic, and these repeats were proposed as the sites of the interaction between α2δ-1 and TSPs (Eroglu et al., 2009). Here we found that a second interaction that is mediated by the third Type I (Properdin-like) domain of TSP1 is responsible for conferring its cell type-specific synaptogenic activity.

β1-Integrin was previously studied for its roles in synaptic plasticity and transmission in hippocampus (Chan et al., 2006; Huang et al., 2006; Ning et al., 2013), glycinergic synapse formation in spinal cord neurons (Charrier et al., 2010) and AMPAR clustering in cortical neurons (Fossati et al., 2019). Our study reveals a new function for β1-Integrin in the CNS: providing cell-type specific control of excitatory synapse formation via glia-secreted synaptogenic factor TSP1. In the retina, β1-Integrin is almost exclusively expressed by the RGCs, and is significantly enriched in the CART-positive ooDSGCs and β1-Integrin is necessary and sufficient for TSP1-mediated synaptogenic activity in the RGCs. Our findings also show that for TSP1 to be synaptogenic it needs to interact both with α2δ-1 and with β1-Integrin, indicating that these two TSP1 receptors work together in neurons to control synaptogenic signaling. In summary, our findings reveal a cross-talk between β1-Integrin and α2δ-1-signaling during synaptogenesis. Future studies investigating the biochemical underpinnings of this crosstalk would be fruitful in elucidating synaptogenic mechanisms in the CNS.

### Lack of TSP1 impairs direction-tuning function of ooDSGCs

Many signaling mechanisms and developmental events shape the formation of DS circuits and their constitutive cell types in the retina. For example, Sema6A, is necessary for symmetric SAC arborization (Sun et al., 2013). Similarly, Cadherins mediate homophilic or heterophilic interactions to recruit the dendrites of distinct RGCs and axons of BCs to form the organized IPL scaffold (Duan et al., 2014; Duan et al., 2018). Also, a transcription regulator Satb1, controls bi-stratification of the ooDSGC arbors via regulating the expression of a cell adhesion molecule, Contactin 5 (Peng et al., 2017). MEGF10 is necessary for the mosaic organization of SACs and the proper targeting of SAC dendrites (Ray et al., 2018). Collectively, these studies illustrate that the proper dendritic arborization and lamination is key to DS circuit formation, and anatomical disruption of these circuit elements frequently results in diminished DS circuit function (Ray et al., 2018).

Our study shows that MG-neuron signaling is also required for proper DS circuit formation and function. We found that the direction selective responses of the ooDSGCs remain in TSP1 KO retinas; however, DS tuning function of the circuit was significantly perturbed as shown by significantly increased tuning width and decreased tuning strength (Figure 5). There were no arborization changes or overt structural deficits found in IPL organization in TSP1 KO retinas, indicating that impaired synapse development caused by a lack of TSP1 is the major contributing factor to the DS deficits observed in the KOs.

The direction-tuning in these circuits is strongly influenced by inhibition (Demb, 2007; Poleg-Polsky and Diamond, 2016). We found that in TSP1 KO retinas, the diminished excitatory synapses in the DS sublayers of the IPL is concurrent with a reduction in the number of inhibitory synapses throughout the IPL. This raises the possibility that TSPs may also shape inhibitory synapse formation. Previous studies implicated TSP1 and β1-Integrin in inhibitory (glycinergic) synapse regulation in the spinal cord (Charrier et al., 2010; Hennekinne et al., 2013). However, others have shown that TSP1 does not induce formation of inhibitory synapses in glia-depleted hippocampal neuron cultures (Hughes et al., 2010). In our in vitro experiments using purified RGCs and ACs, we did not find any evidence for TSP1 to induce GABAergic or cholinergic synapses. Hence, it is plausible that the reduction of inhibitory synapses in the TSP1 KO retinas is driven by a consequent compensatory mechanism induced by the loss of normal excitatory connectivity.

### Does TSP1 serve as a glial cue that temporally and spatially instructs synapse formation?

In the brain, TSP1 and TSP2 are primarily expressed by astrocytes during early postnatal development (P7-P10), coinciding with the initiation of rapid synapse formation, but are not abundant in the mature brain (Cahoy et al., 2008; Christopherson et al., 2005). However, a recent study showed that chemogenetic activation of Gi pathway in striatal astrocytes results in a robust increase in TSP1 transcription and a subsequent rapid excitatory synaptogenesis in nearby neurons, suggesting that this protein could be utilized as a local synaptogenic cue that is released by glia in response to changes in neuronal activity (Nagai et al., 2019). In the rodent retina, the expression levels of TSP1 and TSP2 and their synaptic receptor α2δ-1 sharply increases between P14 and P30 (Koh et al., 2018), during the time window following eye-opening and coinciding with rapid synaptogenesis (Tian, 2008). However, different than brain astrocytes, MG produce TSP1 and TSP2 and deposit them in retinal synaptic layers throughout late postnatal development and well into adulthood (Koh et al., 2018). TSP1 protein is particularly enriched within the DS-sublayers (Koh et al., 2018) suggesting a function for TSP1 at these synapses. Concordantly, in TSP1 KO mice the excitatory synapses in DS layers of the IPL were significantly decreased, despite the fact that these animals still have TSP2 protein, which functionally can compensate for TSP1 *in vitro*. Moreover, MG-specific overexpression of SD1, the synaptogenic TSP1 fragment, was sufficient to promote synapse formation in both WT and KO retinas and the synaptogenic effects of overexpressed SD1 were seen throughout the entire IPL. These results show that glia-mediated TSP1 signal is necessary and sufficient for excitatory synapse development of the DS circuit in the retina and suggest that localized production and/or secretion of TSP1 by MG onto ooDSGC dendrites in the IPL is necessary to form proper DS-circuit connections. Future studies investigating how MG target TSP1 production and/or secretion at the IPL sublaminae containing DS circuit connections would provide further insight into the potential pathways through which glia regulates assembly and function of specialized circuits.

## Methods

### Lead Contact and Materials Availability

Further information and requests for resources and reagents should be directed to and will be fulfilled by the Lead Contact, Cagla Eroglu.

### Experimental Model and Subject Details

#### Animals

All experiments were conducted in accordance with the institutional animal care and use committee guidelines (IACUC Protocol Numbers A-147-17-06 and A-126-18-05). B6.129S2-Thbs1tm1Hyn/J (TSP1 KO), B6;129-Itgb1tm1Efu/J (Itgb1 flox), B6.Cg-Gt(ROSA)26Sortm14(CAG-tdTomato)Hze/J (Ai14), B6.Cg-*Cdh6^tm1.1(cre/ERT2)Jrs^*/J (Cdh6-CreER) were purchased from Jackson laboratory. For RGCs and ACs primary culture, Sprague-Dawley rats with P6 pups were purchased from Charles river. The RGCs and ACs were purified from P7 pups. To achieve conditional KO of Itgb1 in ooDSGCs, we crossed the Itgb1 flox mice to Cdh6-CreER for inducible Cre expression and Ai14 Cre reporter line.

#### Isolation and Culture of Primary Rat Retinal Ganglion Cells (RGCs) and Amacrine Cells (ACs)

RGCs were purified by sequential immuno-panning from P7 (postnatal day 7) Sprague-Dawley rat retinas (Charles River) as previously described (Barres et al., 1988; Goldberg et al., 2002). Retinas were dissected and dissociated with papain (6 U/mL, Worthington). Dissociated cells were panned three times with Bandeiraea Simplicifolia Lectin I (BSL, Vector laboratories) coated Petri dishes to remove immune cells, cell debris and fibroblasts. The unbound cells were transferred to a Petri dish coated with anti-Thy1 (clone T11D7) antibody for specific isolation of RGCs. Floating cells were subsequently transferred to a Petri dish coated with Vc1.1 antibody for amacrine cells. The purified RGCs and ACs were gently trypsinized and re-plated onto poly-D-Lysine (PDL) and laminin-coated glass coverslips in 24-well plates. For RGCs only culture, 35,000 cells/coverslip were plated, and for co-culture of RGCs/ACs, 15,000 RGCs and 60,000 ACs were plated to a coverslip. RGCs and ACs were cultured in a serum-free growth medium containing B27 (1X, Invitrogen), brain-derived neurotrophic factor (BDNF, 50 ng/mL), ciliary neurotrophic factor (CNTF, 10 ng/mL), insulin (1X, GIBCO) and forskolin (5 ng/mL).

#### Retina Preparation for Immunohistochemistry

The retinas from TSP1 WT and KO littermate pairs or Itgb1 WT (0/0) and Itgb1 cKO (f/f) littermate pairs of either sex were collected at P30. For the IHC analyses of TSP1 WT and KO pairs, TSP1 Het (+/-) breeding pairs were utilized, then littermates pair of WT (+/+) and KO (-/-) were collected together. Animals were terminated at P30 by perfusing with Tris-Buffered Saline (TBS, 25 mM Tris-base, 135 mM NaCl, 3 mM KCl, pH 7.6) supplemented with 7.5 µM heparin and followed by 4% paraformaldehyde (PFA). Eyes were removed and immersed in 4% PFA for two hours and subsequently infiltrated with 30% sucrose overnight at 4°C. Eyes were embedded in OCT (frozen tissue matrix) and cut in sequence (12 µm horizontal sections apart) on a cryostat. Every tenth section was placed on the same slide as the first section and a total of four sections (120 µm apart) were collected per genotype per slide. The retina from a littermate KO was sectioned in the same manner and placed on the same slides with the WT.

For Itgb1 cKO, Itgb1(f/+) RTM(Tg/Tg) mice were bred to Itgb1(f/+) Cadherin6 Cre-ERT2 (Tg/Tg) to achieve Itgb1 (+/+) and Itgb1 (f/f) with same copy number of RTM (Tg/0) and Cre (Tg/0). To achieve the Cre-recombination in the ooDSGCs and SACs, tamoxifen was administered (20 mg/mL, 0.6 mg total) subcutaneously twice at P9 and P10. Earlier injection of tamoxifen resulted in heterogenous recombination result shown by tdTomato expression in BC and other ACs.

#### Quantification of Synapses in the Retina

3-4 animals per genotype of TSP1 WT and KO or Itgb1 WT and cKO were used for synapse analysis. Three independent retina sections per each group were used for immunohistochemistry. 5 µm thick confocal z-stacks (optical section depth 0.33 µm, 15 sections/z-stack) of the retinas were imaged at 60× magnification on a Olympus Fluoview confocal laser-scanning microscope. Maximum projections of three consecutive optical sections (corresponding to 1 µm total depth) were generated from the original z-stack. Analyses were performed blind as to genotype. The Puncta Analyzer plugin (written by Barry Wark, modified by Chaichontat (Richard) Sriworarat, available upon request from Cagla Eroglu at) for either ImageJ (NIH; http://imagej.nih.gov/ij/) or FIJI (https://imagej.net/Fiji/Download) was used to count the number of co-localized puncta. This quantification method is based on the fact that pre-and post-synaptic proteins (such as VGluT1 and PSD95) are not within the same cellular compartments of neurons (axons versus dendrites, respectively) and would only appear to partially co-localized at synaptic junctions due to their close proximity. This quantification method yields an accurate estimation of the number of synapses both in vitro and in vivo, because it measures co-localization as opposed to staining of a single pre-or postsynaptic protein that often accumulate in extrasynaptic regions during the course of their life cycle. In agreement, numerous previous studies by ourselves and others have shown that synaptic changes observed by this quantification method is verified by techniques such as electron microscopy and electrophysiology (Allen et al., 2012; Christopherson et al., 2005; Eroglu et al., 2009; Koh et al., 2015; Kucukdereli et al., 2011; Risher et al., 2014; Singh et al., 2016). Details of the quantification method have been described previously (Ippolito and Eroglu, 2010). Briefly, 1 µm thick maximum projections are separated into red and green channels, backgrounds are subtracted (rolling ball radius = 50), and thresholds are determined in order to detect discrete puncta without introducing noise. Minimum pixel size of puncta was set as 4 to remove any background noise. The Puncta Analyzer plugin then uses an algorithm to detect the number of puncta that are in close proximity across the two channels, yielding quantified co-localized puncta. In order to calculate percentage of WT co-localization, co-localized puncta values for WT were averaged, then all image values (WT and KO) were normalized to the ratio of the calculated WT average.

#### Quantification of Synapses in the cultured RGCs

To test the RGC-subtype specific synaptogenic function of TSPs, 35,000 RGCs were plated on glass coverslips, and then recombinant human TSP1 or TSP2 (R&D) were added at DIV4 and DIV7. To block TSP-induced synapse formation, 32 uM Gabapentin (GBP) was added to the TSPs. On 10 DIV, RGCs were fixed with 4% PFA (w/v) and stained for pre-and post-synaptic markers Bassoon (mouse anti-bassoon, 1:1000) and Homer-1 (rabbit anti-homer, 1:500), or Basoon, PSD95 (biotinylated anti-PSD95, 1:250) and CART (rabbit anti-CART, 1:1000). For detection Alexa-conjugated secondary antibodies (Invitrogen, goat anti rabbit AF-488 (1:500) and goat anti mouse AF-594 (1:500), or goat anti mouse IgG2a AF-647 (1:250), Streptavidin AF-568 (1:250) and goat anti rabbit AF-488 (1:1000)) were used for detection. Coverslips were mounted in Vectashield mounting medium with DAPI (Vector Laboratories) on glass slides (VWR Scientific). RGCs were imaged on a Zeiss Axioimager M1 Epifluorescence Microscope (Carl Zeiss) using a 63X oil objective. Morphologically healthy single cells that were at least two cell diameters from their nearest neighbor were identified randomly by DAPI fluorescence. At least 25-30 cells per condition (CART positive or negative) per treatment were imaged and analyzed per experiment. The results presented are the average of 2-3 independent experiments. Captured images were analyzed for co-localized synaptic puncta with a custom Image J plug-in described above.

#### RGC Transfections

RGCs were transfected using the Lipofectamine LTX (Invitrogen) reagent at DIV3. Briefly, 250μl conditioned culture medium was transferred from the cells saved in another plate after combining with 250 μl of fresh culture medium at 37°C in 10% CO_2_ incubator. The RGCs were then fed with 200μl fresh media. The DNA/Lipofectamine mix was made as per manufacturer’s protocol with 1:5 DNA to LTX ratio. A total of 250 ng DNA was transfected per well of 24-well plate. Briefly, 125 ng of each DNA construct expressing BFP and ITGB1 (Addgene 51920) and 1.25μl of Lipofectamine LTX was used in each well (in a 24 well plate). After 2 hours, RGCs were washed with DPBS twice and were fed with previously saved 500 uL fresh and conditioned culture medium. Treatment of purified synaptogenic domain fragments (SD1 or SD2, 150 ng/mL) was started on the next day (DIV4). RGCs were stained for synapses after 6 days of treatment as described above in the synapse assay section. The number of synapses on transfected RGCs identified by BFP expression was quantified as described above.

#### Immunohistochemistry

Retina sections were washed three times then permeabilized in PBS with 0.3% Triton-X 100 (PBST; Roche, Switzerland) at room temperature. Sections were blocked in 5% Normal Goat Serum (NGS) or Normal Donkey Serum (NDS) in PBST for 1 hr at room temperature. Primary antibodies (mouse anti-Bassoon 1:500 [RRID: AB_10618753, ADI-VAM-PS003-F, Enzo, NY], guinea pig anti-VGlut1 1:750 [AB5905, Millipore, MA], rabbit anti-PSD95 1:500 [RRID: AB_87705, 51–6900, Invitrogen, CA], mouse anti-Gephyrin 1:250 [RRID: AB_1279448, 147-021, Synaptic Systems, Goettingen, Germany], mouse anti-Glutamine Synthetase 1:1,000 [RRID: AB_397879, 610517, BD Biosciences, CA], goat anti-Choline Acetyltransferase [RRID: AB_11214092, AB144P, Millipore], rabbit anti-Homer1 [RRID: AB_887730, 160-002, Synaptic Systems], rabbit anti-CART [RRID: AB_2313614, H00362, Phoenix Pharmaceutic], biotinylated anti-PSD95 [RRID: AB_2619799,124-011BT, Synaptic Systems], rat anti-tdTomato 16D7 clone [RRID: AB_2732803, EST203, Kerafast], guinea pig anti-RBPMS [RRID:AB_2687403, ABN1376, EMD Millipore], guinea pig anti-VAChT [RRID: AB_10893979, 139-105, Synaptic Systems] were diluted in 5% NGS or 5% NDS containing PBST. Sections were incubated overnight at 4°C with primary antibodies. Secondary Alexa-fluorophore (488, 568, 594 and 647) conjugated antibodies (Invitrogen) were added (1:200 in PBST with 5% NGS or 5% NDS) for 2 hr at room temperature. Slides were mounted in Vectashield with DAPI (Vector Laboratories, CA) and images were acquired on an Olympus Fluoview confocal microscopy using 60X oil lens.

#### Retina Preparation for Multi-electrode Array (MEA) Recording

Retinas from two pairs of littermate P30 WT and TSP1 KO mice with BL6/J background were used for the MEA recordings. All mice were housed in a 12 hr light-dark cycle with access to food and water. Animals were dark adapted (12 hrs) prior to the euthanasia then dissection of the retinas was performed in a dark room with the help of infra-red light (>900 nm) and infra-red goggles to minimize photobleaching of the retina. The eyes were removed and transferred to a petridish containing sodium buffered Ames’ solution (Sigma) with pH equilibrated to 7.4 by bubbling 95% O2 and 5% CO2. The eyes were hemisected and vitreal attachment to the retina was removed. A piece of dorsal retina (1-2 mm) was isolated by detaching it from the pigment epithelium. Dorsal side was identified by previously established vasculature landmarks (Wei et al., 2010), then placed to the MEA with ganglion cell side down. The MEA is consisted of 519 electrodes with 30 um spacing, with 450um on each side (Gunning et al., 2007). Extracellular voltages were digitized at 20 kHz and stored for post-hoc analysis. Spikes were identified by applying a voltage threshold to each electrode. Principal component analysis of identified waveforms yielded a subspace with clusters of spikes which were isolated using a mixture of Gaussian models. Each spike cluster was identified as a putative RGC if the refractory period exceeded 1.5 ms with <10% contamination with at least 100 spikes. Duplicate clusters identified from temporal cross-correlation were removed, leaving a subset of RGC spike trains which were used for further analysis. For further details on spike sorting, see (Field et al., 2007; Litke, 2004).

#### Visual Stimulation

Stimuli were created in MATLAB (Mathworks, Natick, MA), displayed on a gamma-corrected OLED with 60.35 Hz refresh rate, and focused on the photoreceptor outer segment through the microscope objective (Eclipse Ti-E 4x, Nikon, Nikon Corporation). The mean intensity of the stimulus on the photoreceptors was ∼7000 photoisomerizations/rod/sec, or ∼5000 photoisomerizations/cone/sec for M-conopsins.

Two types of stimuli were used: (1) Moving bars (at 50% Weber contrast) traveling in twelve different directions at four different speeds ranging from 480 µm/sec to 1920 µm/sec. The movement in each direction, for each speed, was repeated eight times. (2) Checkerboard binary patterns with 40 x 40 µm^2 checker size, refreshing at 60.35 Hz.

#### ooDSGC Classification and Quantification of Direction Tuning Function

To distinguish DSGCs from non-DSGCs, spiking responses of individual RGCs to bars moving in different directions were used to calculate direction selective index (DSI):

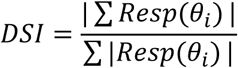

where *Resp* (*θ_i_*) is the response to a bar moving along the direction *θ_i_*. The DSI values for all RGCs at two speeds: 480 µm/sec and 1448 µm/sec, for all RGCs were plotted on a log-log axis, and the cluster of RGCs with DSI values higher than 0.5 were selected as DSGCs. For all recordings, a DSI threshold of 0.5 provided clear separation between clusters of DSGCs and non-DSGCs (Figure 3B). From within the DSGC cluster, ooDSGCs were identified based on prominence of ON and OFF peaks in PSTHs in the preferred direction at speeds 480 µm/sec and 1448 µm/sec (Yao et al., 2018). ON DSGCs with relatively weak OFF peaks were excluded in subsequent analyses.

To estimate the width of tuning curve, first the number of spikes elicited for each movement direction was calculated. This yielded a magnitude of normalized vector sum of spikes: R, that lies between 0 and 1. Mapping the normalized responses in each direction to the circular Normal distribution (Oesch et al., 2005), the circular standard deviation σ*_circ_* is obtained as:

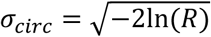

To measure the tuning strength, the difference between responses in the preferred and null directions were normalized by their sum. Because directions were sampled at 30° interval, response in the preferred direction (PD) or null direction (ND) was calculated by taking a cosine weighted average of responses to the two neighboring movement directions around the preferred or null directions. This yielded the following equation for tuning strength:

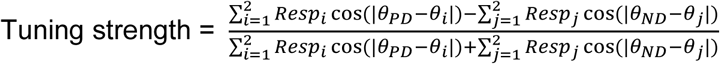

A tuning strength value of 1 implies response in the null direction is zero, while a tuning strength value of 0 implies responses in the preferred and null directions are equal.

#### RNA Fluorescence In Situ Hybridization (FISH)

We visualized mRNA for Itgb1 using the RNAcope (ACD Bio, Newark, CA). A set of FISH probes targeting Itgb1 (514281, RNAscope Probe - Mm-Itgb1) was purchased from Advanced Cell Diagnotics. Another probe set targeting a bacterial gene DapB (310043, RNAscope Negative Control Probe-DapB) was used as a negative control. Cryo-sectioned retina samples prepared as described above were used for probing RNA molecules. Probe hybridization and signal amplification was performed as manufacturer’s protocol following ‘RNAscope® Fluorescent Multiplex Kit User Manual’. The sections were dehydrated and permeablized with series of 50%, 70% and 100% ethanol for 5 mins each at room temperature. After wash and rehydration with PBS, the sections were incubated with RNA probes for 2 hrs at 40°C in a humidified chamber. The hybridized probes were amplified following sequential washes and incubation procedures with the RNAscope detection kit per manufacturer’s protocol. Probes were visualized by Atto 550 fluorophore included in the RNAscope detection kit. The immunohistochemistry was performed after FISH procedures to visualize RGC cell bodies (anti-RBPMS) and CART-positive cell bodies (anti-CART) by following the IHC protocol as decribed above omitting the detergent, Triton-X. Detection of the RNA probes and IHC signal was performed using the Olympus Fluoview confocal microscopy with 60X oil lens.

#### His-tagged protein purification

For production and purification of the chimeric synaptogenic domain fragments, the gene block fragments that transcribe SD2-T1*, SD2-T2*, SD2-T1*T2* and SD1-T1^ were synthesized (IDT) than inserted into a vector backbone pAPtag5 using XbaI and NheI enzyme sites. Strong signal peptide IgK leader sequence was inserted in the N-terminal each gene block to promote secretion of the translated peptides and 6X His-tag was added on the C-terminal of each fragment. The final plasmid DNA was collected using the Endofree Maxi-prep kit (Qiagen). To produce each fragment, total of five 10 cm dishes were used per peptide production and 10 ug of DNA was transfected to a 10 cm dish of HEK cells (8 × 10^6^ per 10 cm dish). The transfected HEK cells were conditioned for 3 to 4 days before collection of the medium.

For His-tag purification, the conditioned medium was collected then concentrated using the Vivaspin column (MWCO 5K). The concentrated medium was mixed with Ni-NTA resin (Qiagen) then incubated at 4C overnight while gently nutating. Next day, the resin was washed with DPBS then collected in a Poly-Prep Chromatography column (Thermo Fisher). The peptide fragments were eluted with 250 mM Imidazole in DPBS. The elutes were run on the SDS-PAGE gel and the gel was stained with coomassie blue to confirm the purity. Concentration was further assessed using the micro BCA protein assay kit (Thermo) per manufacturer’s protocol.

#### Adeno-Associated Virus (AAV) Production and Intravitreal AAV Injection

For production of AAV expressing SD1 and/or mCherry, Plasmids AAV-gfaABC1D-SD1-IRES-mCherry and AAV-gfaABC1D-IRES-mCherry were used to produce the AAV particles for the overexpression and control, respectively. AAV was produced by transfecting these plasmids (30 µg) to the HEK cells with helper (pAD-delta F6, 15 µg) and the capsid plasmids (pXX2-ShH10Y445F, 15 µg, Pellissier et al., 2014) specific for the Muller glia using the Polyethylenimine (PEI, 1.25 mM). A total of five 15 cm tissue culture dishes (12 × 10^6^ HEK cells per dish) were used to produce one type of virus. Three days post-transfection, HEK cells were lysed using the cell lysis buffer (Cell lysis buffer: Add 3 ml of 5 M NaCl and 5 ml of 1 M Tris-HCl (pH 8.5) to 80 ml of dH2O. Adjust the pH to 8.5 with NaOH and adjust the volume to 100 ml with dH2O. Sterilize by passing through a 0.22-μm filter and store at 4 °C.). Then the AAV particles were released from the cells by three cycles of freeze/thaw between dry ice-ethanol and 37 °C water bath. Then AAV were purified using Iodixanol Gradient Ultracentrifugation as described by Addgene (https://www.addgene.org/protocols/aav-purification-iodixanol-gradient-ultracentrifugation/). Briefly, the cell lysates are loaded to the gradients of iodixanol (15%, 25%, 40% and 60%) in the OptiSeal tubes (Beckman Coulter, Indianapolis, IN). Then the lysates were spin down at 67,000 rpm for 1 hour at 18 °C with the ultracentrifuge (Beckman). The AAV particles were collected by aspirating the interface between 40-60% iodixanol with a syringe. The collected AAV-containing solution was mixed with ice-cold DPBS and concentrated using the Vivaspin column (100 MWCO) at 4°C. Collected virus particles were aliquoted and stored at −80 °C until use.

TSP1 WT or KO mouse pups at postnatal day 5 were used for intravitreal injections. The retinas from AAV injected animals were collected at P30 as described above. A total of 2 µL containing ∼1×10^12^ vg/mL of AAV was injected intravitreally to transduce the MG using 5 µL Hamilton syringes. Transduction to the MG was identified by the expression of mCherry as described in Figure 4.

#### Statistical Analysis

Statistical analyses of the quantified synapse number data were performed using Student’s t-test or one-way analysis of variance (ANOVA) followed by post-hoc test (Tukey’s HSD), if applicable. JMP Genomics Pro 14.0 software (SAS, Cary, NC) was used for all statistical analysis of the data. All data was expressed as mean±SEM, and significance was demonstrated as * *p*<0.05, *** p<0.0001.

## Acknowledgements

Parts of this work were supported by grants from the National Institutes of Health (NEI 1F32 EY027997 and NIA 2T32AG000029 to S.K., R01 EY024567 and R01 EY027193 to G.F and EY024694 to J.N.K.), a NEI Core Grant for Vision Research (P30; EY5722) to Duke University, Duke University Chancellor’s Award to C.E., Whitehead Scholars Program to G.F., Ruth K. Broad Postdoctoral Fellowship to S.R. and Duke Regeneration Next Initiative Postdoctoral Fellowship to S.K..

## Author Contributions

S.K., S.R., J.K., G.F. and C.E. designed experiment, analyzed data and wrote the paper. S.K., S.R., O.E., S.S. and W.J.C., collected and analyzed data.

