## Supplementary Figures for "Thrombospondin-1 Promotes Circuit-Specific Synapse Formation via β1-Integrin"

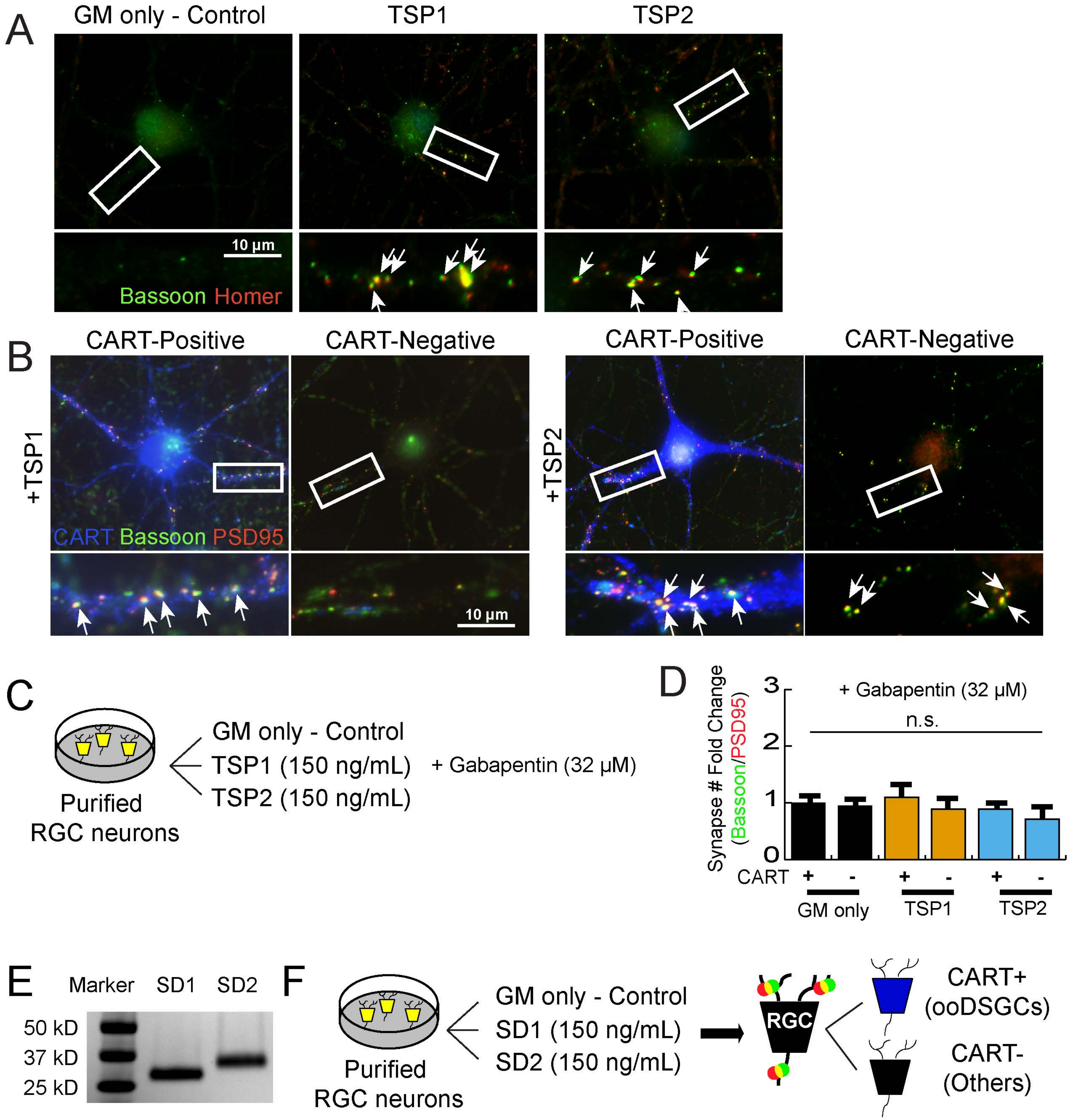
A
GM only - Control
TSP1
TSP2
10 µm
Bassoon Homer
B
CART-Positive
CART-Negative
CART-Positive
CART-Negative
+TSP1
+TSP2
CART Bassoon PSD95
10 µm
C
GM only - Control
TSP1 (150 ng/mL)
TSP2 (150 ng/mL)
+ Gabapentin (32 µM)
Purified
RGC neurons
D
+ Gabapentin (32 µM)
n.s.
Synapse # Fold Change
(Bassoon/PSD95)
0
1
2
3
CART
+
-
+
-
+
-
GM only
TSP1
TSP2
E
Marker
SD1
SD2
50 kD
37 kD
25 kD
F
Purified
RGC neurons
GM only - Control
SD1 (150 ng/mL)
SD2 (150 ng/mL)
RGC
CART+
(ooDSGCs)
CART-
(Others)

**Supplementary Figure 1. TSP1 promotes synapse formation in CART-positive subpopulation of RGCs. Related to Figure 1. (A)** Representative images of RGC dendrites from cells treated with growth medium only (GM only control) or TSP1 (150 ng/mL) or TSP2 (150 ng/mL). The RGCs are immuno-stained with antibodies specific for presynaptic (Bassoon, green) and postsynaptic (Homer, red) proteins. The inlets (white boxes) are shown in higher magnification and the co-localized synaptic puncta (merge, yellow) are marked with white arrows. **(B)** Representative images of RGC s treated with TSP1 (150 ng/mL) or TSP2 (150 ng/mL). ooDSGCs are labeled with CART (blue) in addition to the synaptic markers, Bassoon (pre-, green) and PSD95 (post-, red). The inlets (white boxes) are shown in higher magnification and the co-localized synaptic puncta (merge, yellow) are marked with white arrows. **(C)** Schematic representation of the experimental design. RGCs are isolated from P7 retinas then treated with purified synaptogenic domain fragments of TSP1 or TSP2 in the presence of Gabapentin (32 μM). **(D)** Both TSP1 and TSP2 induced synaptogenic function is inhibited in the presence of the small molecule inhibitor, Gabapentin (32 μM). **(E)** SDS-PAGE gel analysis of purified synaptogenic domain fragments, SD1 and SD2. (500 ng/lane). **(F)** Schematic representation of the experimental design for the Figure 1H-J. RGCs are isolated from P7 retinas then treated with purified synaptogenic domain fragments of TSP1 (SD1) or TSP2 (SD2).

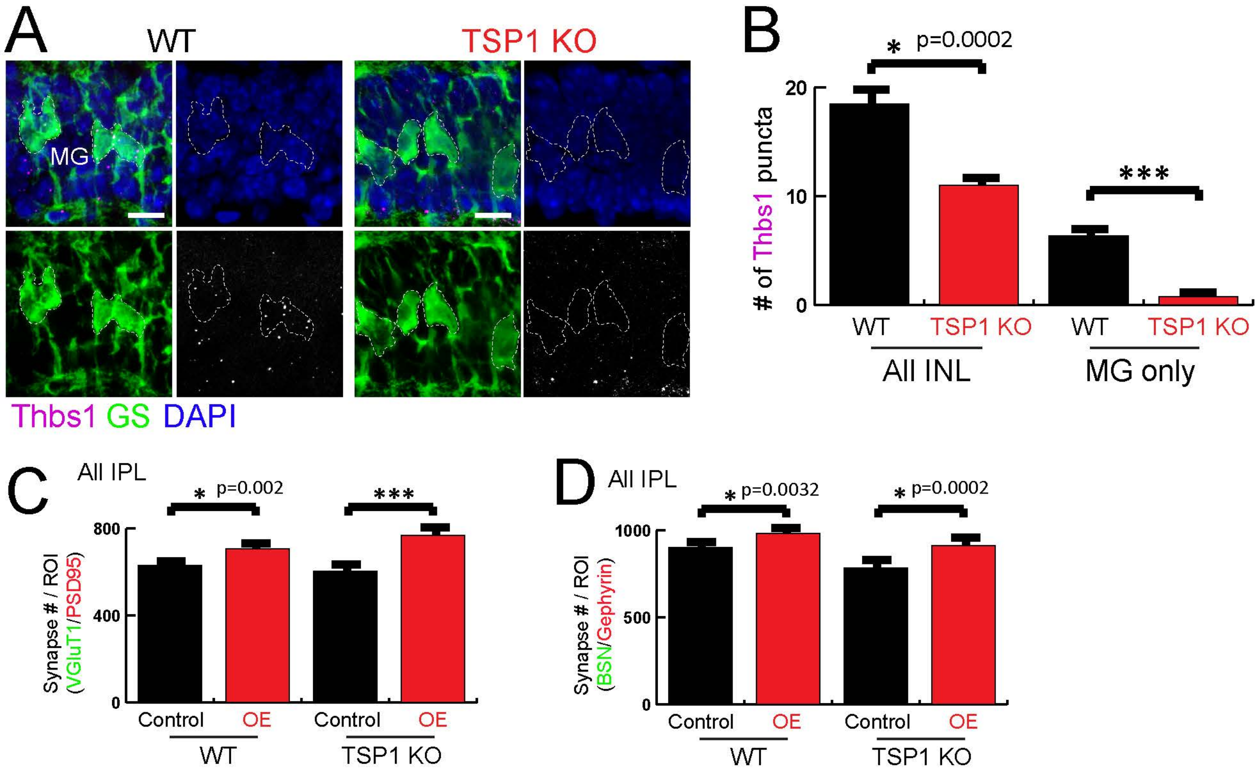
A
WT
TSP1 KO
MG
Thbs1 GS DAPI
B
* p=0.0002
***
### of Thbs1 puncta
20
10
0
WT
TSP1 KO
WT
TSP1 KO
All INL
MG only
C
All IPL
* p=0.002
***
Synapse # / ROI
(VGluT1/PSD95)
800
400
0
Control
OE
Control
OE
WT
TSP1 KO
D
All IPL
* p=0.0032
* p=0.0002
Synapse # / ROI
(BSN/Gephyrin)
1000
500
0
Control
OE
Control
OE
WT
TSP1 KO

**Supplementary Figure 2. MG-produced TSP1 is sufficient to induce synapse formation in DS circuit IPL sublayers. Related to Figure 4. (A)** Representative images of mRNA probes for Thbs1 (magenta) in the mouse P30 retina. MG are labeled by anti-GS (green). **(B)** Quantification of Thbs1 mRNA probes found in all INL or MG cell bodies demonstrates exclusive expression of Thbs1 in MG cell bodies. **(C and D)** Over-expression (OE) of SD1 induced excitatory and inhibitory synapse formation in the IPL in both wild-type and TSP1 KO (n=3-4 mice per genotype per treatment, One-way ANOVA, *** $p<0.0001$).

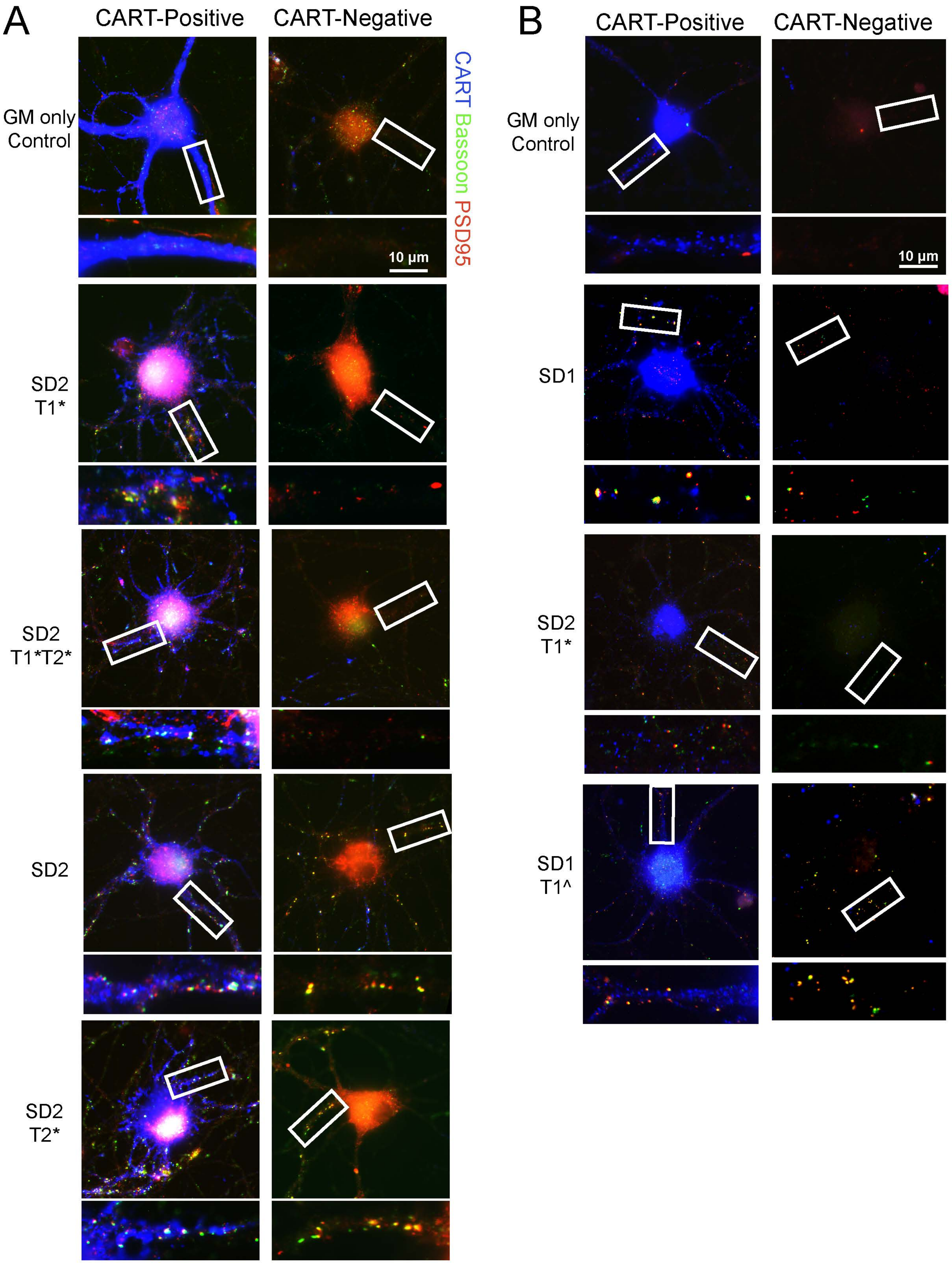
A
CART-Positive
CART-Negative
GM only Control
SD2 T1*
SD2 T1*T2*
SD2
SD2 T2*
CART Bassoon PSD95
10 μm
B
CART-Positive
CART-Negative
GM only Control
SD1
SD2 T1*
SD1 T1^
CART Bassoon PSD95
10 μm

**Supplementary Figure 3. Type 1 repeat domains of TSP1 and TSP2 confers RGC subtype specificity. Related to Figure 6. (A and B)** Representative images of RGCs and dendrites from cells treated with growth media (RGC alone) only or different synaptic fragments at 150 ng/mL concentration. The RGCs are stained with antibodies specific for presynaptic (Bassoon, green), postsynaptic (PSD95, red) proteins and OODSGCs are stained with CART (Blue). The inlets (white boxes) are shown in higher magnification and the co-localized synaptic puncta (merge, yellow) are marked with white arrows.

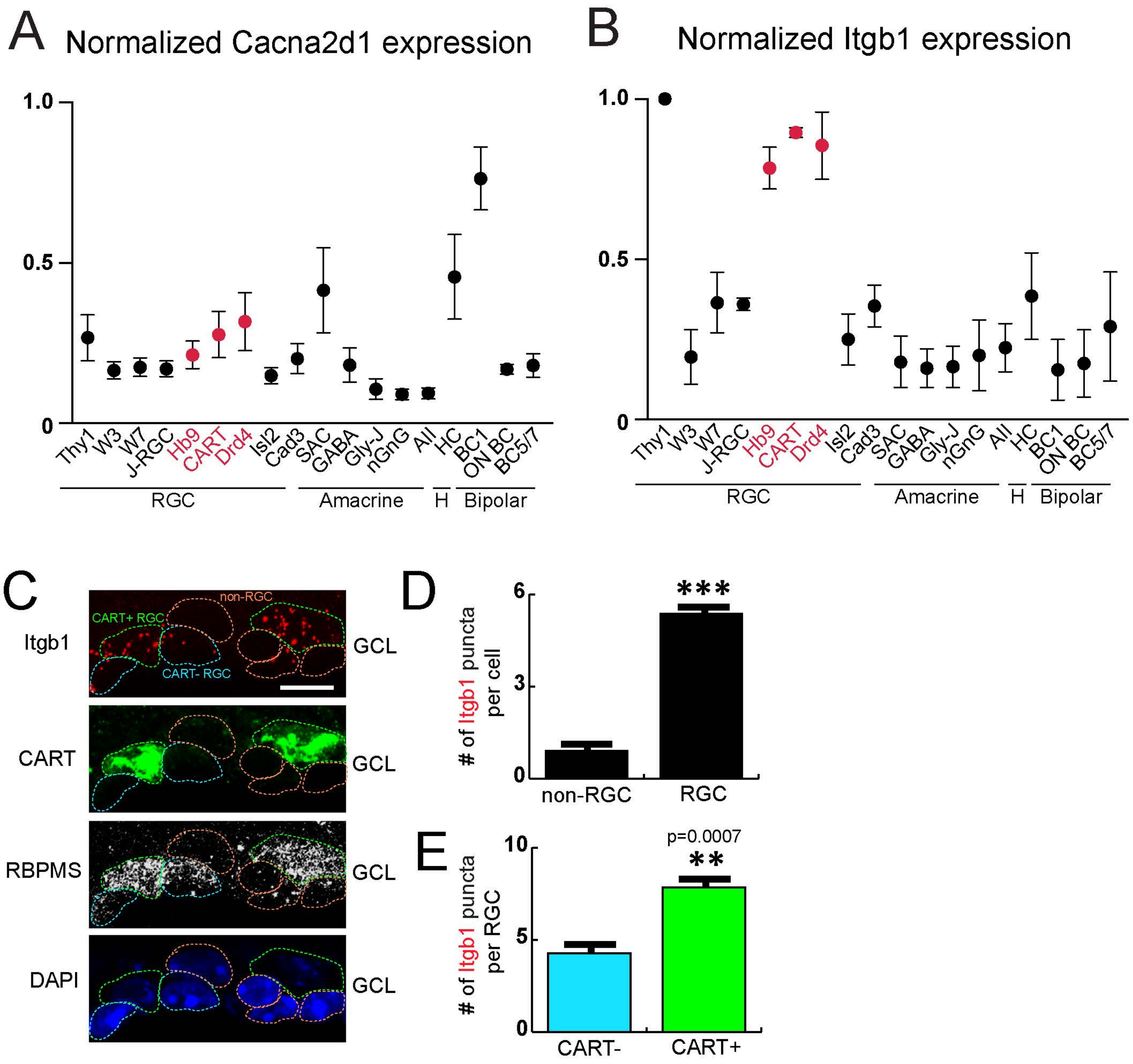
A
Normalized Cacna2d1 expression
1.0
0.5
0
Thy1
W3
W7
J-RGC
Hb9
CART
Drd4
Isl2
Cad3
SAC
GABA
Gly-J
nGnG
AII
HC
BC1
ON BC
BC5/7
RGC
Amacrine
H
Bipolar
B
Normalized Itgb1 expression
C
non-RGC
CART+ RGC
CART- RGC
Itgb1
CART
RBPMS
DAPI
GCL
D
### of Itgb1 puncta per cell
***
non-RGC
RGC
E
### of Itgb1 puncta per RGC
p=0.0007
**
CART-
CART+

**Supplementary Figure 4. Type 1 repeat domains of TSP1 and TSP2 confers RGC subtype specificity. Related to Figure 6.** Normalized relative expression levels of **(A)** Cacna2d1 or **(B)** Itgb1 in RGCs, Amacrine cells, Horizontal Cells (H) and Bipolar cells. The graph is re-plotted from previously reported microarray analyses (Kay et al., 2012). **(C)** Representative images of mRNA probes for Itgb1 (red) in the mouse retina. RGCs are labeled by anti-RBPMS (white) and ooDSGCs are labeled by CART (green). **(D)** Quantification of number of Itgb1 mRNA probes found in pan-RGC (RBPMS-positive) versus non-RGC (RBPMS-negative) demonstrates significant enrichment of Itgb1 expression in RGCs in the retina. **(E)** Quantification of number of Itgb1 mRNA probes found in CART-negative versus –positive RGCs demonstrates enriched expression of Itgb1 in CART-positive RGCs.

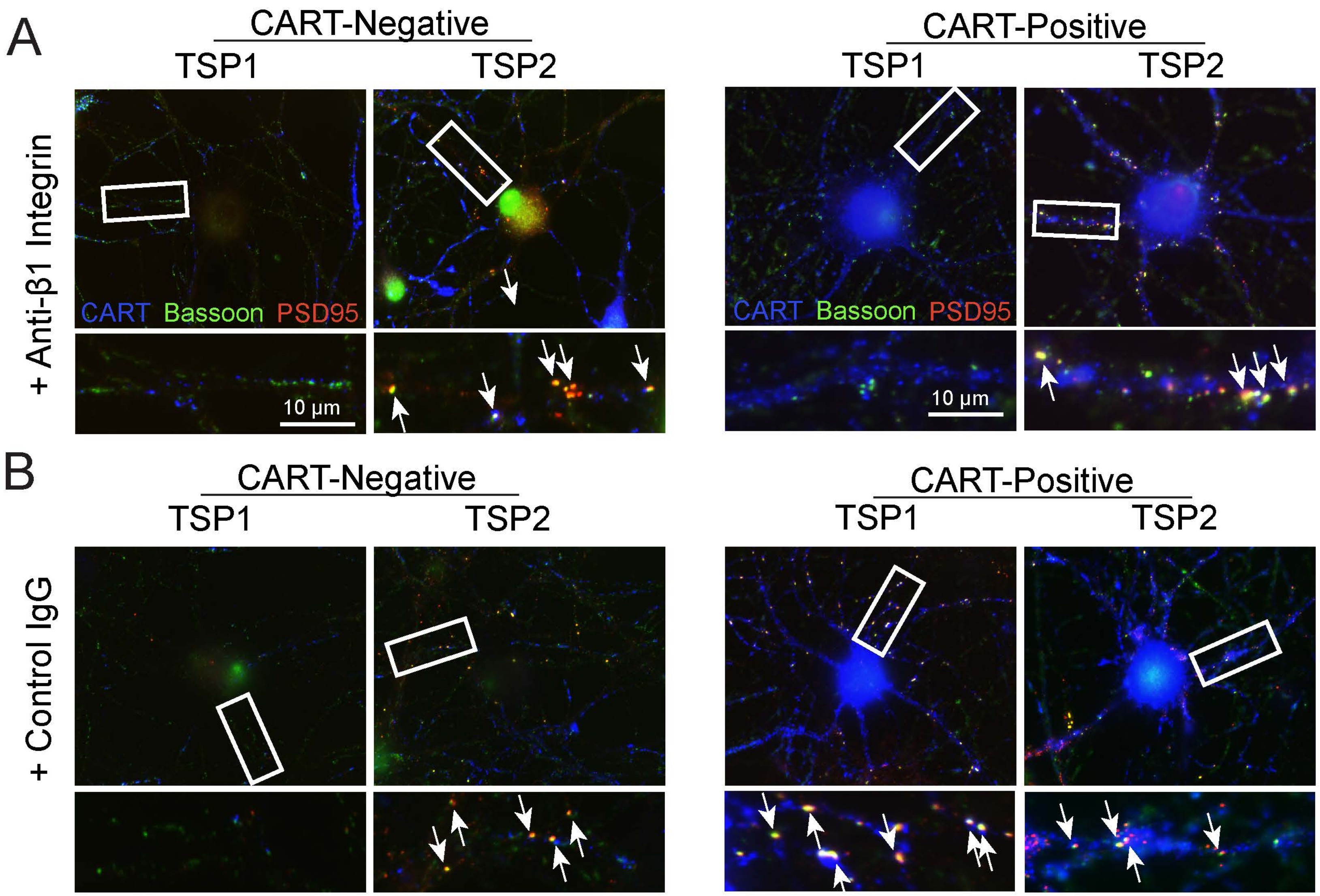
A
CART-Negative
TSP1
TSP2
CART-Positive
TSP1
TSP2
+ Anti-β1 Integrin
CART Bassoon PSD95
10 μm
CART Bassoon PSD95
10 μm
B
CART-Negative
TSP1
TSP2
CART-Positive
TSP1
TSP2
+ Control IgG

**Supplementary Figure 5. ooDSGC-specific synaptogenic activity of TSP1 is mediated via Integrin β1. Related to Figure 7. (A)** Representative images of RGCs and dendrites from cells treated with full-length TSP1 or TSP2 in the presence of Integrin β1 function blocking antibody or **(B)** control IgG. The RGCs are stained with antibodies specific for presynaptic (Bassoon, green), postsynaptic (PSD95, red) proteins and OODSGCs are stained with CART (Blue). The inlets (white boxes) are shown in higher magnification and the co-localized synaptic puncta (merge, yellow) are marked with white arrows.

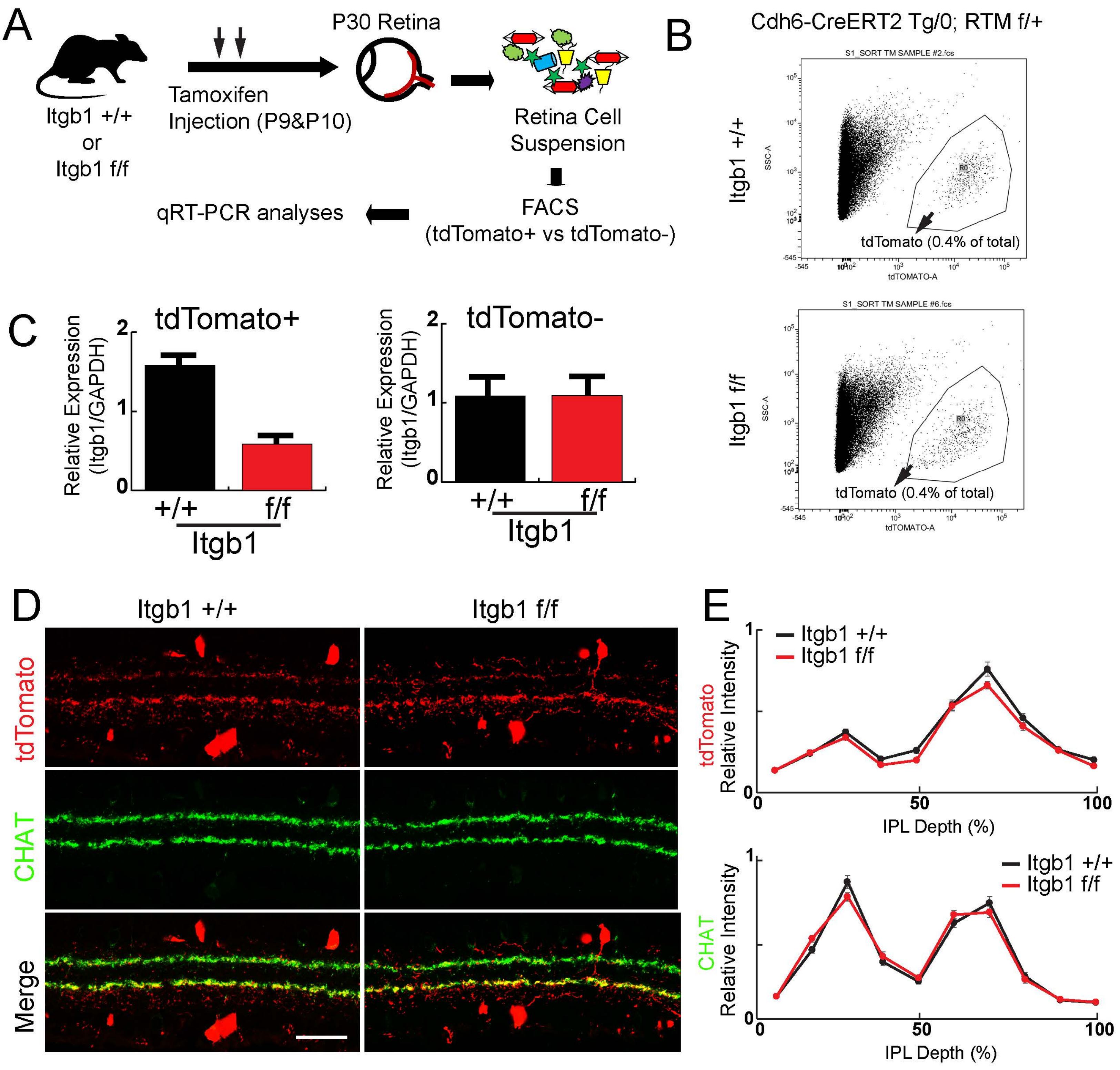
A
Itgb1 +/+
or
Itgb1 f/f
Tamoxifen
Injection (P9&P10)
P30 Retina
Retina Cell
Suspension
FACS
(tdTomato+ vs tdTomato-)
qRT-PCR analyses
B
Cdh6-CreERT2 Tg/0; RTM f/+
Itgb1 +/+
Itgb1 f/f
S1_SORT TM SAMPLE #2.fcs
S1_SORT TM SAMPLE #6.fcs
SSC-A
tdTOMATO-A
R0
tdTomato (0.4% of total)
C
tdTomato+
tdTomato-
Relative Expression
(Itgb1/GAPDH)
+/+
f/f
Itgb1
D
Itgb1 +/+
Itgb1 f/f
tdTomato
CHAT
Merge
E
Itgb1 +/+
Itgb1 f/f
tdTomato
CHAT
Relative Intensity
IPL Depth (%)

**Supplementary Figure 6. Integrin β1 is required for the DS circuit synapse development. Related to Figure 8. (A)** Schematic work flow for isolating td-Tomato/Cre+ cells by fluorescence activated cell sorting (FACS). A single cell suspension from P30 retinas are prepared then passed through a BD FACS sorter to collect tdTomato/Cre-positive or tdTomato/Cre-negative cells. RNA was isolated from these cells, reverse transcribed to cDNA, then the expression level of Itgb1 mRNA was analyzed by qRT-PCR. **(B)** Representative contour plots of control (Itgb1 +/+) and cKO (Itgb1 f/f) FACS analysis. About 0.4% of total sorted cells were tdTomato-positive in both control and cKO. **(C)** Quantitative real-time RT-PCR analysis of Itgb1 expression in tdTomato-positive (left panel) and tdTomato-negative (right panel) cells. Expression level of Itgb1 is reduced by 70% in cKO animals. The expression level was normalized to the expression level of GAPDH using the $2^{-\Delta\Delta CT}$ method. **(D)** Representative images of the tdTomato (red) positive cells and dendrites in WT and Itgb1 cKO (Itgb1 f/f) retinas. DS circuit IPL sublayers are visualized by CHAT (green). **(E)** Relative fluorescence intensity plots of tdTomato- (top) or CHAT-positive (bottom) dendrite staining across IPL between WT and Itgb1 cKO (Itgb1 f/f) retinas.
